# Transport of endo-β-mannosidase AtEBM from cytoplasm to the vacuole through autophagy pathway in *Arabidopsis*

**DOI:** 10.64898/2025.12.09.693153

**Authors:** Huan Wei, Jinyan Mai, Jianyu Su, Gengnan Fang, Shiting Feng, Mantao Chen, Jinxia Yu, Xiao Huang, Na Luo, Faqiang Li

**Author notes:** Corresponding authors: Drs. Na Luo, and Faqiang Li. These authors contributed equally to this work (Huan Wei, Jinyan Mai). The authors responsible for distribution of materials integral to the findings presented in this article in accordance with the policy described in the Instructions for Authors are: Na Luo, and Faqiang Li. **Email addresses :** Huan Wei, Jinyan Mai, Jianyu Su, Gengnan Fang, Shiting Feng, Mantao Chen, Jinxia Yu, Xiao Huang, Na Luo, Faqiang Li.

## Abstract

Autophagy is a conserved eukaryotic intracellular degradation pathway that exhibits specialized forms, such as the biosynthetic cytoplasm-to-vacuole targeting (Cvt) pathway in yeast. This pathway selectively delivers several hydrolase precursors to the vacuole. While the core autophagy machinery is highly conserved between yeast and plants, the existence of a similar Cvt-like pathway for vacuolar hydrolase trafficking in plants remains unclear. Here, we found that *Arabidopsis thaliana* endo-β-mannosidase (AtEBM), a glycosidase involved in degrading vacuolar *N*-glycans released from glycoproteins, is selectively transported to the vacuole via autophagy. Further analyses revealed that AtEBM directly interacts with the core autophagy protein ATG8 through an ATG8-interacting motif (AIM). This interaction is essential for sequestering AtEBM into autophagosomes. Remarkably, the AIM is conserved among EBM orthologs across the green plant lineage. In addition, several phenotypic studies showed that *atebm* mutants generated through CRISPR-Cas9 editing did not exhibit autophagy defects, including early senescence and hypersensitivity to nutrient starvation, but rather had reduced fecundity. Thus, these findings reveal a Cvt-like pathway in plants that fulfills a biosynthetic role by selectively delivering the glycosidase AtEBM to the vacuole.

## INTRODUCTION

Autophagy is an intracellular degradation pathway that is highly conserved among eukaryotes, ranging from yeast to mammals and plants. During this process, cytoplasmic constituents, including soluble and aggregated proteins and organelles, are engulfed by double-membrane structures called autophagosomes. These structures then deliver the constituents to the vacuole in yeast and plants or the lysosome in mammals for degradation (Nakatogawa, 2020; Petersen *et al*., 2024; Zhuang *et al*., 2024). Originally discovered as a response to nutrient starvation, autophagy was long considered as a non-selective, bulk degradation pathway that provided building blocks and energy for survival under nutrient-limiting conditions. However, research over the past two decades has revealed that autophagy is highly selective and tightly regulated. Autophagic receptors can specifically recognize and recruit deleterious protein aggregates, dysfunctional organelles, and invading pathogens into autophagosomes for degradation (Stephani & Dagdas, 2020; Lamark & Johansen, 2021).

In addition to its role as an intracellular degradation pathway, autophagy in budding yeast (*Saccharomyces cerevisiae*) can also selectively transport several vacuolar hydrolases, including aminopeptidase I (ScApe1), aspartyl aminopeptidase (ScApe4), leucine aminopeptidase 3 (ScLap3), and α-mannosidase (ScAms1) from cytoplasm to the vacuole. This unique form of selective autophagy is known as the cytoplasm-to-vacuole targeting (Cvt) pathway, which is biosynthetic rather than degradative (Lynch-Day & Klionsky, 2010; Yamasaki & Noda, 2017). After synthesis, the Cvt cargo ScApeI rapidly forms dodecamers that further assembles into a large complex termed the Cvt complex, together with other cargoes such as ScAms1 (Kim *et al*., 1997). A specific autophagy receptor, autophagy-related protein 19 (ScAtg19), recognizes and recruits the complex to the autophagy machinery, promoting the formation of a special type of autophagosome called the Cvt vesicle. This kind of vesicle is about 150 nm in diameter and much smaller than starvation-induced autophagosomes (300–900 nm in diameter) (Scott *et al*., 2001; Shintani *et al*., 2002).

In budding yeast, approximately 20 autophagy-related (Atg) proteins are essential for the formation of autophagosomes during bulk autophagy. These proteins can be categorized into several functional complexes/groups, including the Atg1 kinase complex (Atg1-Atg13-Atg17-Atg29-Atg31), Atg9-containing vesicles, the Atg2–Atg18 complex, the class III phosphoinositide 3-kinase (PI3K) complex I (Vps34-Atg6–Atg14–Vps15) and the Atg8 and Atg12 conjugation systems (Itakura & Mizushima, 2010). All of these functional complexes are also required for Cvt vesicle formation, although the components of the Atg1 complex differ greatly from those in bulk autophagy; ScAtg11 and ScAtg17 function as protein scaffolds that are essential for selective autophagy and bulk autophagy, respectively (Lynch-Day & Klionsky, 2010; Yamasaki & Noda, 2017). During Cvt vesicle formation, ScAtg11 associates with ScAtg19, which acts as an adapter to recruit the Cvt complex to the phagophore assembly site (PAS), promoting the formation of Cvt vesicles (Shintani *et al*., 2002). Furthermore, lipidated ScAtg8, which is anchored in the inner membrane of Cvt vesicles, was found to tightly interact with ScAtg19. ScAtg19 then tethers Cvt complexes to the phagophore, ensuring their selectively engulfed (Noda *et al*., 2008; Abert *et al*., 2016). In the fission yeast *Schizosaccharomyces pombe*, SpNBR1, an evolutionarily related cargo receptor to ScAtg19, was found to mediate the vacuolar targeting of several cytosolic hydrolases, including Ams1, Lap2, Ape2 and Ape4 (Liu *et al*., 2015; Wang *et al*., 2021). Interestingly, this SpNBR1-mediated vacuolar targeting pathway does not require core Atg proteins, but rather employs the endosomal sorting complex required for transport (ESCRT) to sequester the cargoes into the multivesicular bodies (MVBs) and deliver them to the vacuole (Liu *et al*., 2015).

The lytic vacuole in plants plays a role similar to that of yeast vacuole and the lysosome in mammals. It is involved in the degradation of proteins and other macromolecules, as well as nutrient storage and recycling and detoxification (Shimada *et al*., 2018). Multiple proteomic studies of *Arabidopsis* and other plant species have revealed that, like the yeast vacuole, the lytic vacuole in plants is home to numerous transporters and hydrolases (Carter *et al*., 2004; Jaquinod *et al*., 2007; Ohnishi *et al*., 2018). Studies over the last several decades have shown that the vacuolar trafficking routes and components required for the targeting of these transporters and hydrolases are partially conserved between plants and yeast. However, plants have also evolved specific variations in the organization of their vacuolar trafficking systems and the protein machinery that regulates trafficking (Aniento *et al*., 2022). The best-characterized route is the one depending on the sequential action of small RAB GTPases RAB5 and RAB7, which is similar to the classical carboxypeptidase (CPY) pathway in yeast (Ebine *et al*., 2014; Hecht *et al*., 2014). Vacuolar proteins targeted by this pathway, such as barley (*Hordeum vulgare*) aleurain, generally have signal sequences that direct them to the endoplasmic reticulum (ER) for synthesis and then to the Golgi apparatus. After that, the proteins are sorted into the trans-Golgi network (TGN)/early endosome (EE) and transported to the vacuole via MVBs. A second route also involves the TGN and MVBs and requires only RAB5 for delivering several tonoplast proteins, such as SYNTAXIN OF PLANTS22 (Ebine *et al*., 2014) and the vacuolar H^+^ ATPase subunit VHA-a3 (Feng *et al*., 2017). A third pathway utilizes the AP-3 adaptor to transport cargoes directly from the Golgi, which is similar to the yeast alkaline phosphatase (ALP) pathway (Feraru *et al*., 2010; Zwiewka *et al*., 2011; Ebine *et al*., 2014). Additionally, there is mounting evidence pointing to the existence of a Golgi-independent ER-vacuole trafficking pathway for unconventional vacuolar cargoes (Park *et al*., 2004; Isayenkov *et al*., 2011; Teper-Bamnolker *et al*., 2021).

Glycoprotein turnover is essential for maintaining cellular homeostasis. Most glycoproteins are degraded by proteases in the vacuole/lysosome, while the *N*-glycan moieties attached to the asparagine residues of glycoproteins are released and degraded by a series of vacuolar/lysosomal glycosidases, as well as cytosolic glycosidases (Winchester, 2005; Harada *et al*., 2015). Although the complete picture of plant N-glycan degradation is not yet understood, a unique set of vacuolar glycosidases has been shown to be involved in the degradation process (Figure 1A, Léonard *et al*., 2009; Ishimizu, 2015; Kato *et al*., 2018). The major plant N-glycans M5A (high mannose-type) and M3FX (plant complex-type) are transported from cytosol to the vacuole and degraded cooperatively by at least four glycosidases: α-mannosidase, endo-β-mannosidase (EBM), β1,2-xylosidase, and α1,3-fucosidase.

**Figure 1.**
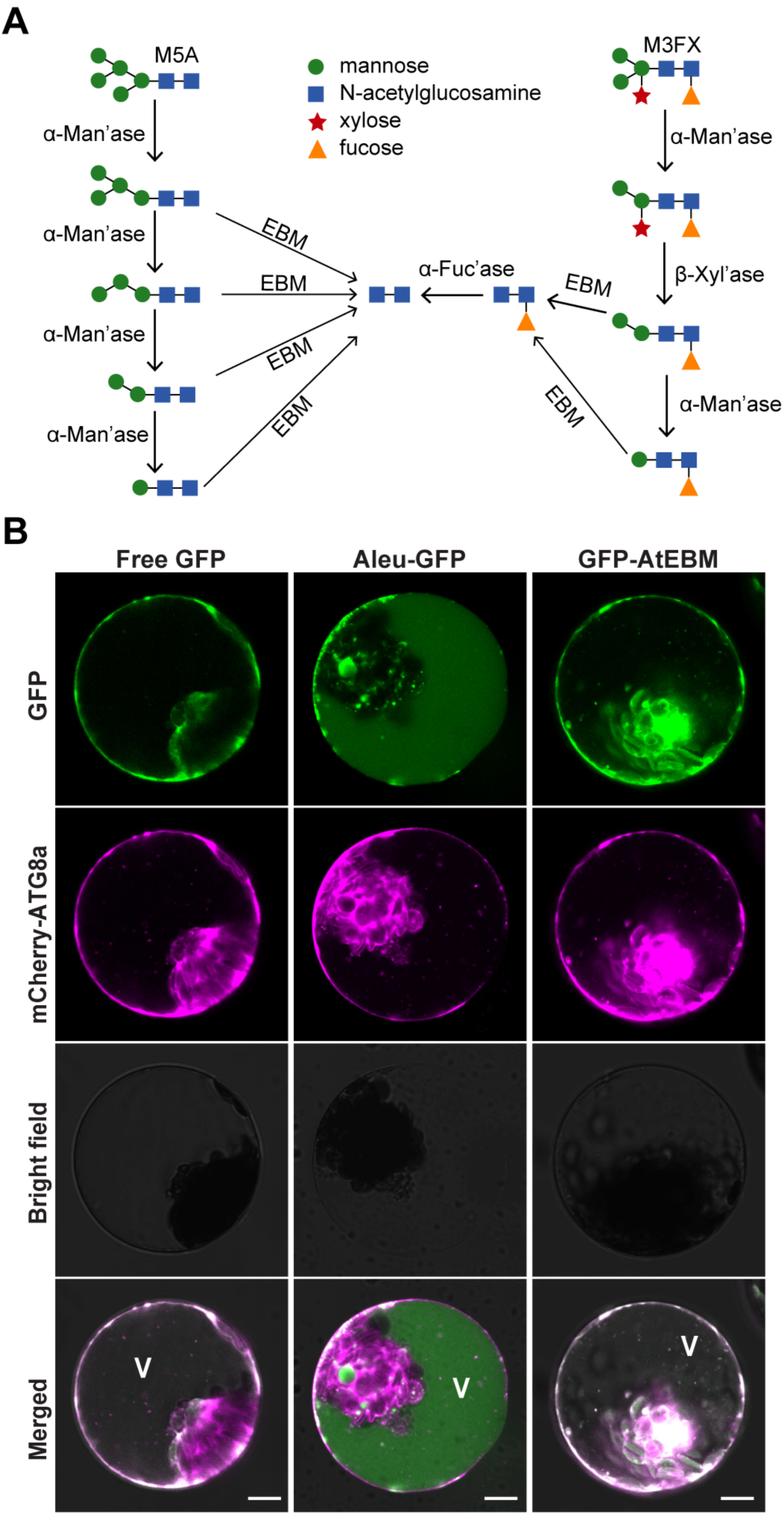
Subcellular localization of *Arabidopsis N*-glycan degrading enzyme AtEBM in protoplasts. **(A)** The proposed vacuolar degradation pathway of plant *N*-glycans. M5A and M3FX are the major high mannose-type and plant complex-type *N*-glycans in plant cells, respectively. In *Arabidopsis*, the degradation of these glycans involves at least four glycosidases: α-mannosidase (α-Man’ase), endo-β-mannosidase (EBM), β1,2-xylosidase (β-Xyl’ase), and α1,3- fucosidase (α-Fuc’ase). **(B)** Subcellular localization of AtEBM in *Arabidopsis* protoplasts. Protoplasts were co-transformed with the autophagy marker mCherry-AtATG8a and plasmids expressing the indicated gene constructs, then analyzed with confocal microscopy 12 to 14 hours after transfection. Non-fused GFP protein was used as a cytosolic control, and Aleu-GFP was used as a lytic vacuole marker control. Concanamycin A (ConA), an inhibitor of vacuolar-type ATPases, was used to block vacuolar degradation. V, vacuole. Bars = 10 μm.

Although the autophagy machinery is highly conserved in eukaryotic cells, whether the existence of a Cvt-like pathway in plants for targeting vacuolar hydrolases remains unclear. Here, we demonstrate that *Arabidopsi*s endo-β-mannosidase (AtEBM) is selectively transported to the vacuole via autophagy. Further analysis revealed that the direct interaction of AtEBM with the core autophagy protein ATG8 is essential for the sequestration of AtEBM into autophagosomes. Thus, we propose that plants use a Cvt-like pathway to fulfill a biosynthetic function by delivering AtEBM selectively to the vacuole.

## METHODS

### Plasmid construction

To generate the constructs for transient expression in *Arabidopsis* protoplasts, the full-length cDNAs of *EBM* were amplified from the leaf tissues of *Arabidopsis* and rice (*Oryza sativa*), and from the vegetative tissue of liverwort (*Marchantia polymorpha*) using the gene-specific primers listed in Supplemental Table S1, and inserted into the pBI221-GFP vector. To generate the mCherry-AtNBR1 construct, the full-length cDNA of *AtNBR1* was amplified and placed into the pBI221-mcherry vector. The lytic vacuole marker Aleu-GFP, the autophagy markers GFP-AtATG8a and mCherry-AtATG8a, the autophagy cargo receptor GFP-AtNBR1 and the late endosome mCherry-RABG3f was previously reported (Dong *et al*., 2025). The Golgi marker Man1-RFP, the TGN marker RFP-SYP61, the MVB/PVC marker mCherry-Rha1, and the tonoplast marker mCherry-VAMP711 were obtained from Professor Liwen Jiang’s laboratory. The ER marker CNX-mCherry was obtained from Professor Caiji Gao’s laboratory.

To generate the constructs for investigating the protein-protein interaction between AtEBM and AtATG8a using luciferase (LUC) complementation imaging (LCI) assay, the *AtEBM* and *AtATG8* CDSs were amplified and cloned into the pCAMBIA1300-nLUC and pCAMBIA1300-cLUC vector, respectively (Chen *et al*., 2008). The AtEBM-nLUC vector containing a mutated ATG8-interacting motif (AIM) was generated by PCR mutagenesis. The cLUC-AtATG8a vectors containing wild-type AtATG8a, mutated LDS variant [cLUC-ATG8a(ΔLDS)] or mutated UDS [cLUC-ATG8a(ΔUDS)] were described previously (Dong *et al*., 2025). To investigate the conservation of the EBM-ATG8 interaction, the full-length EBMs and ATG8s were amplified from cDNAs prepared from rice and liverwort, and cloned into the pCAMBIA1300-nLUC and pCAMBIA1300-cLUC vectors, respectively. The EBM-nLUC vectors containing the mutated AIM were generated by PCR mutagenesis.

The constructs for the yeast two-hybrid assay were generated by cloning the CDSs of *AtEBM*, *AtATG8a* and *AtNBR1* into the pGBKT7 and pGADT7 vectors (Clontech, USA). Constructs containing AtEBM^mAIM^, AtATG8a(ΔLDS), and AtATG8a(ΔUDS) were generated by PCR mutagenesis.

### Plant materials and growth conditions

The *A. thaliana* ecotype Columbia-0 (Col-0) was used as the wild type for this study. The T-DNA mutants *atg5-1* (Thompson *et al*., 2005), *atg7-2* (Chung *et al*., 2010), *atg9-3* (Shin *et al*., 2014), *atg11-1* (Li *et al*., 2014) and *nbr1-2* (Zhou *et al*., 2013) were described previously. All *Arabidopsis* seeds were surface-sterilized using gaseous chlorine and stratified by soaking in distilled water at 4°C for two days. The seeds were spread on full-strength Murashige and Skoog (MS) medium at 22°C under a long-day condition (LD, 16 h light/8 h dark) for seven days. Thereafter, the seedlings were transferred to soil and grown until harvested under LD conditions. Tobacco (*Nicotiana benthamiana*) seeds were sown directly in the soil for germination. Two weeks later, seedlings were transferred to individual pots and grown at 25°C under LD conditions.

To produce GUS transgenic plants, the 2,000-bp region upstream of the putative translation start site of *AtEBM* was amplified by PCR and inserted into the pCAMBIA1305 vector between the *Hind*III and *Spe*I sites to fuse with the GUS reporter. The construct, which was verified by sequencing, was then introduced into the wild-type Col-0. Multiple independent transgenic lines were obtained by hygromycin-based selection, and two T3 homozygous lines were chosen for further study.

To generate transgenic *Arabidopsis* plants overexpressing GFP-AtEBM, the full-length *AtEBM* cDNA was cloned into the vector pCAMBIA1300 vector between the *Xma*I and *Kpn*I restriction sites under the control of *UBQ10* promoter, to create the *proUBQ10::GFP-AtEBM* vector. To generate the *proAtEBM::GFP-AtEBM* construct, the *AtEBM* native promoter was amplified from the *proAtEBM::GUS* vector and replaced the *UBQ10* promoter in the *proUBQ10::GFP-AtEBM* plasmid to generate the *proAtEBM::GFP-AtEBM* vector. After sequencing verification, the constructs were delivered into the *Agrobacterium tumefaciens* strain GV3101 and transformed into the Col-0 wild-type using the floral dip method. Homozygous plants expressing GFP-AtEBM were obtained in the T3 generation through hygromycin-based selection and fluorescence microscopy observation. The *GFP-AtEBM atg7-2* and *GFP-AtEBM mCherry-ATG8f* plants were generated by crossing the *proUBQ10::GFP-AtEBM* transgenic plants with the *atg7-2* mutant and the *mCherry-ATG8f* transgenic lines (Zhuang *et al*., 2013), respectively.

To develop *atebm* mutants, two CRISPR/Cas9 vectors, each expressing two guide RNAs, were constructed using an egg cell-specific promoter-controlled CRISPR/Cas9 genome editing system (Xing *et al*., 2014; Wang *et al*., 2015). Guide RNAs that target *AtEBM* were designed using the CRISPR-GE website (http://skl.scau.edu.cn/) and cloned into the pHEE401E vector with primers listed in Supplemental Table S1. The sequencing-verified constructs were then introduced into the wild-type Col-0 using the Agrobacterium-mediated floral-dip method. The resulting T1 transformants were selected on MS medium supplemented with 50 ng/µL hygromycin and were then genotyped via PCR amplification and sequencing of the genomic regions encompassing the DNA target sites.

### Plant treatments and phenotyping

The autophagy-related phenotypic analyses were carried out as previously described by Huang *et al*. (2019). Briefly, plants were grown on soil under short-day conditions (12 h light/12 h dark) at 22°C for eight weeks to detect the early senescence phenotype. For nitrogen (N) starvation treatments, seeds were germinated and grown vertically on solid MS medium with or without N for seven days, after which the root lengths were measured and quantified. Alternatively, seven-day-old seedlings grown on liquid MS medium with N were transferred to fresh MS medium with or without N for an additional week, after which the chlorophyll content was determined and quantified. For carbon starvation, two-week-old seedlings grown on solid MS medium without sucrose (Suc) were wrapped in aluminum foil and kept in continuous darkness for 12 to 14 days. After the dark treatment, the stressed plants were allowed to recover under LD conditions for 12 days, after which the survival rates of seedlings were quantified.

The *in vitro* and *in vivo* pollen germination assays were performed as previously described (Dong *et al*., 2025). For silique length measurement, the plants were grown at 22°C under LD conditions and the 5^th^-12^th^ siliques on the main inflorescence were selected. The siliques were then divided into three categories based on their length: fully fertile (Type I, >10 mm), partially sterile (Type II, 5–10 mm), and completely sterile (Type III, <5 mm).

### Protein-protein interaction assays

Protein-protein interactions by Y2H were performed using the Matchmaker™ Gold Yeast Two-Hybrid System (Clontech, USA) according to the manufacturer’s instructions. The pairwise combination of genes in the pGBKT7 and pGADT7 vectors were co-transformed into the yeast strain AH109, with empty vectors serving as the negative controls. The presence or absence of protein-protein interactions was assessed based on the growth of yeast on synthetic dextrose (SD) minimal medium lacking Ade, His, Trp, and Leu (SD-Ade-His-Trp-Leu).

Protein-protein interactions in *planta* were determined using the LCI assay as described (Chen *et al*., 2008). In brief, various combinations of nLUC and cLUC constructs, as well as empty vectors, were introduced into the *A. tumefaciens* strain GV3101 and then co-infiltrated into tobacco leaves. Protein-protein interactions were determined based on the luciferase activity, which was measured using the NightSHADE LB 985 system (Berthold Technologies, USA), with firefly D-luciferin applied 36-48 hours after infiltration.

The glutathione S-transferase (GST) pull-down assay was conducted as previously described (Dong *et al*., 2025). In brief, GST-ATG8e was first produced in *Escherichia coli* BL21 (DE3) and purified using Glutathione-Sepharose resins (C650031, Sangon Biotech Co., Ltd., China) according to the manufacturer’s instructions. For the pull-down analysis, total proteins were extracted from c. 1 g of seven-day-old *Arabidopsis* seedlings expressing GFP-AtEBM and mixed with the GST-ATG8e-bound resins in the binding buffer (140 mM NaCl, 2.7 mM KCl, 10 mM Na_2_HPO_4_, 1.8 mM KH_2_PO_4_, pH 7.4). After incubating for one hour at 4°C, the beads were then washed five times with 1 mL of wash buffer (140 mM NaCl, 2.7 mM KCl, 10 mM Na_2_HPO_4_, 1.8 mM KH_2_PO_4_, pH 7.4), each wash lasting one minute. Samples boiled with loading buffer were separated by SDS-PAGE and subjected to immunoblot analysis.

### GUS staining

The GUS assay was performed with *proAtEBM::GUS* transgenic plants. The plant tissues were fixed in 90% acetone at 4°C for 30 minutes, and then washed three times with washing solution (0.1 M sodium phosphate buffer, pH 7.0, 10 mM sodium EDTA, 2 mM potassium ferrocyanide, 2 mM potassium ferricyanide). The samples were then incubated in GUS staining solution (0.1 M sodium phosphate buffer at pH 7.0, 10 mM sodium EDTA, 2 mM potassium ferrocyanide, 2 mM potassium ferricyanide, and 1 mg/mL 5-bromo-4-chloro-3-indolyl-β-glucuronic acid) at 37°C in the dark until distinct blue spots appeared. The reaction was terminated by the addition of 70% anhydrous ethanol. Specimen were then photographed using a Nikon SMZ1270 stereomicroscope.

### Co-expression network analysis

The co-expression data for *AtEBM* were obtained from the publicly available ATTED-II database (version 12; http://atted.jp/). A set of 100 query genes, including *AtEBM*, was used to carry out the co-expression network analysis using Network Drawer tool. The coexpression option was set to ‘add a few genes’, and the PPI option was set to ‘Draw PPIs’. The coexpressed genes were further interpreted by GO functional enrichment analysis to identify the overrepresented functional categories in the network.

### Construction of phylogenetic trees

Protein sequences were aligned using Clustal Omega (http://www.ebi.ac.uk/Tools/msa/clustalo/) Then, the circular maximum likelihood tree was constructed using IQ-TREE (version 2.1.4; Nguyen *et al*., 2015) with the best-fit Q.pfam+R6 substitution model (lowest Bayesian Information Criterion score). Branch support was calculated using 1000 bootstrap replicates and the Shimodaira-Hasegawa-like approximate likelihood ratio test. The final phylogenetic trees were annotated using iTOL v7.1.1 (Letunic & Bork, 2016).

### Transient expression in protoplasts

Transient expression in *Arabidopsis* protoplasts was performed using the PEG-calcium transfection method as described previously (Yoo *et al*., 2007). Mesophyll protoplast cells were prepared from the rosette leaves of four-week-old plants. After PEG-mediated transfection, the protoplasts were incubated at 22°C in the dark for 12 to 16 hours before confocal microscopy observation. To detect autophagic bodies in the vacuolar lumen, 1 μM V-ATPase inhibitor concanamycin A (BVT-0237-M001, AdipoGen Life Sciences, USA) or an equivalent volume of dimethyl sulfoxide (DMSO) was added to the protoplasts and incubated for 12 hours before observation.

### Fluorescence confocal microscopy imaging and analysis

All confocal images were acquired using a Leica Stellaris 5 confocal laser scanning microscope. To observe transgenic plants expressing GFP-AtEBM, six-day-old seedlings grown on MS solid medium were transferred to either fresh MS medium, nitrogen-deficient medium or sucrose-free medium supplemented with 1 μM ConA for the indicated time, and fluorescence images of the root elongation zone were obtained. Observation of the fluorescence in plants coexpressing the dual reporters GFP-AtEBM and mCherry-ATG8f was conducted using the same methods as described above. For GFP observation, the excitation wavelength was 488 nm, and the fluorescence signal was detected at a range of 490-540 nm. To simultaneously acquire both GFP and mCherry signals, excitation wavelengths of 488 nm for GFP and 543 nm for mCherry were used alternatively in the multitrack mode of the microscope with line switching. Quantitative analysis of images was performed by using ImageJ software (https://imagej.net/) as described by Kim *et al* (2022).

### Protein isolation and immunoblot analyses

Total crude proteins were extracted from *Arabidopsis* seedlings by homogenizing them in a 2X SDS-PAGE sample buffer containing 10% (v/v) β-mercaptoethanol. The homogenates were then vortexed for five minutes, boiled at 100°C for five minutes, and clarified by centrifugation at 13000 *g* for ten minutes. The resulting supernatant was then separated by SDS-PAGE and transferred onto polyvinylidene difluoride membranes (IPVH00010; Millipore) for immunoblot analysis. The following specific antibodies were used in the protein blotting analysis: ATG8 (Thompson *et al*., 2005); GFP (HT801-02, TransGen Biotech, China); histone H3 (ab1791, Abcam, UK); GST (B1153, Biodragon, China). The blots were developed using a BeyoECL Plus kit (P0018S, Beyotime Biotech., China), according to the manufacturer’s instructions.

## RESULTS

To identify potential vacuolar hydrolases whose trafficking is mediated by autophagy, we focused on glycosidases and searched for those identified in the *Arabidopsis* vacuolar proteomic samples (Carter *et al*., 2004; Jaquinod *et al*., 2007; Ohnishi *et al*., 2018). These three proteomic studies detected a total of 47 glycosidases belonging to 20 families, eight of which were identified in all three experiments (Table S2). We then used UniProt (https://www.uniprot.org/) to retrieve the detailed annotations of these glycosidases, including molecular weights, isoelectric points, predicted signal peptides, and subcellular locations. 36 of the 47 detected glycosidases were predicted to contain signal peptides or propeptides, suggesting that these glycosidases likely rely on the classic multivesicular body (MVB)-mediated trafficking pathway for vacuolar transport. Of the 11 glycosidases predicted to lack signal peptides, the subcellular localizations of seven proteins were predicted to be outside the vacuole: TRE1 (At4g24040), MNS2 (At3g21160), MNS3 (At1g30000), DPE2 (At2g40840), BAM5 (At4g15210), At3g47000, and At3g54440. In addition, these seven proteins were only detected in one of the proteomic studies, suggesting that they may be nonvacuolar components.

To investigate the subcellular localizations of the remaining four glycosidases [AtEBM (At1g09010), ENGase85A (At5g05460), At3g47040 and At3g47050], we created the N-terminal GFP-tagged constructs of these glycosidases, and transiently coexpressed them with autophagy marker mCherry-ATG8a and analyzed their distribution patterns by confocal microscopy. As shown in Figure 1B, one of the glycosidases, GFP-AtEBM was detected in the vacuolar lumen after treatment with concanamycin A (ConA), a specific V-ATPase inhibitor, and showed strong colocalization with mCherry-ATG8a. This observation is consistent with previous enzymatic studies that showed AtEBM has an acid pH-optimum (Ishimizu *et al*., 2004; Sasaki *et al*., 2005). In contrast, the cytosolic control (free GFP) and the lytic vacuolar marker (Aleu-GFP; Humair *et al*., 2001), a GFP tagged with the vacuolar sorting determinant of barley aleurain, were found mainly in the cytoplasm and vacuolar lumen, respectively. They did not co-localize with mCherry-ATG8a. Unlike AtEBM, the subcellular locations of the other three glycosidases were primarily in the cytosol rather than the vacuolar lumen (data not shown).

The AtEBM is unique in that it lacks a signal peptide and has been detected in all three proteomic studies (Table S2). Furthermore, AtEBM has been identified in the immunoprecipitants of FP-tagged ATG8 in multiple proteomic studies (Zeng *et al*., 2021; Zhao *et al*., 2022; Zhou *et al*., 2023), suggesting its association with autophagy. Therefore, we focused on the vacuolar trafficking of AtEBM. We then used a protoplast transient assay to examine whether the subcellular location of GFP-AtEBM is altered by autophagy mutations defective in various core ATG components. *Arabidopsis* possesses an Atg11 counterpart, which contains structural features of both Atg11 and Atg17 and is essential for both selective and bulk autophagy (Li *et al*., 2014); ATG9 vesicles supply proteins and lipids during autophagosome progression (Zhuang *et al*., 2017), and ATG7 is an E1-like enzyme in the ATG8- and ATG12-conjugation systems (Doelling *et al*., 2002). In contrast to wild-type protoplasts, no GFP-AtEBM puncta were observed in *atg11-1*, *atg9-3* or *atg7-2* mutant cells, regardless of ConA treatment (Figure 2). Taken together, these results suggest that the transport of GFP-AtEBM to the vacuole is mediated by autophagy.

**Figure 2.**
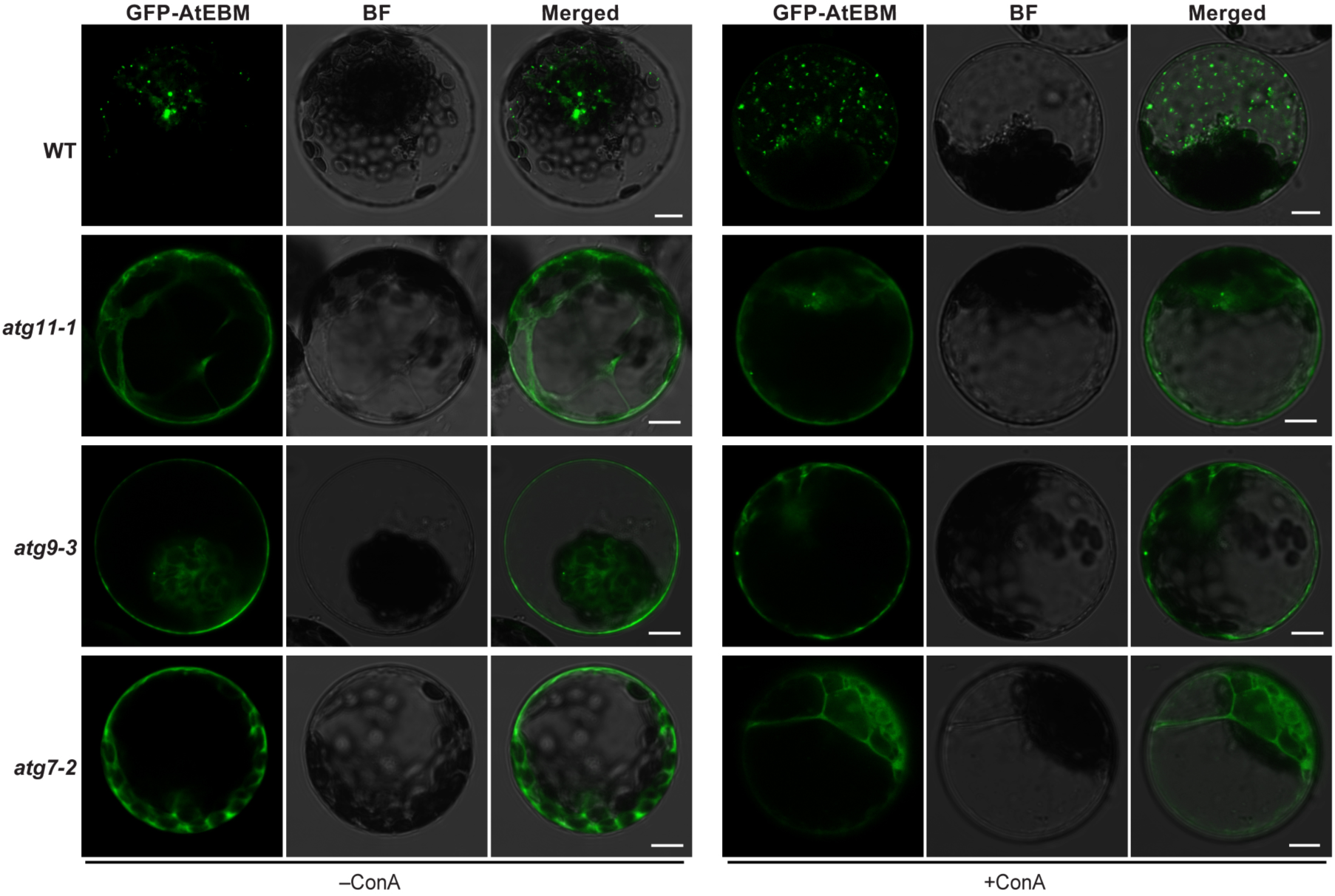
Mutations in core *ATG* components block the vacuolar transport of GFP-AtEBM. Leaf protoplasts from wild type (WT) or various *atg* mutants were transformed with the GFP-AtEBM construct and treated with 1 μM ConA (+ConA) or DMSO (–ConA) for 12 to 14 hours before confocal imaging analysis. Bars = 10 μm.

### NBR1 is not essential for the vacuolar transport of GFP-AtEBM

Previous studies have shown that the Cvt cargoes tend to self-oligomerize and are transported to the vacuole mediated by the receptors Atg19 or its paralog Atg34 in *S. cerevisiae*, and by the receptor Nbr1 and the ESCRT machinery in *S. Pombe* (Liu *et al*., 2015; Yamasaki & Noda, 2017; Wang *et al*., 2021). Although *Arabidopsis* homologs of Atg19 or Atg34 are not apparent, a functional ortholog of the autophagic receptor NBR1 has been described (Svenning *et al*., 2011). Therefore, we attempted to determine whether AtNBR1 is the receptor for the vacuolar trafficking of AtEBM. First, we used the standard yeast two-hybrid (Y2H) assay and the luciferase complementation imaging (LCI) assay to examine the interaction between AtNBR1 and AtEBM. As shown in Figures 3A and 3B, no interaction could be detected between the two proteins. Additionally, no self-interaction of AtEBM was detected using the Y2H and LCI assays, which differs from yeast Cvt cargoes (see Supplemental Figure 1). When GFP-AtEBM and mCherry-AtNBR1 were coexpressed in *Arabidopsis* protoplasts, poor co-localization signals were observed (Figure 3C). Only 8.0% and 12.6% of the GFP-AtEBM-labelled puncta appeared to colocalize with the mCherry-AtNBR1 puncta, with or without ConA treatment, respectively (Figure 3D). *Arabidopsis* protoplast transient expression studies also showed that the vacuolar deposition of GFP-AtEBM was not significantly different between the wild-type control and the *nbr1-2* mutant (Figures 3E and 3F). Taken together, these results indicate that AtNBR1 is not the receptor responsible for targeting AtEBM to the vacuole.

**Figure 3.**
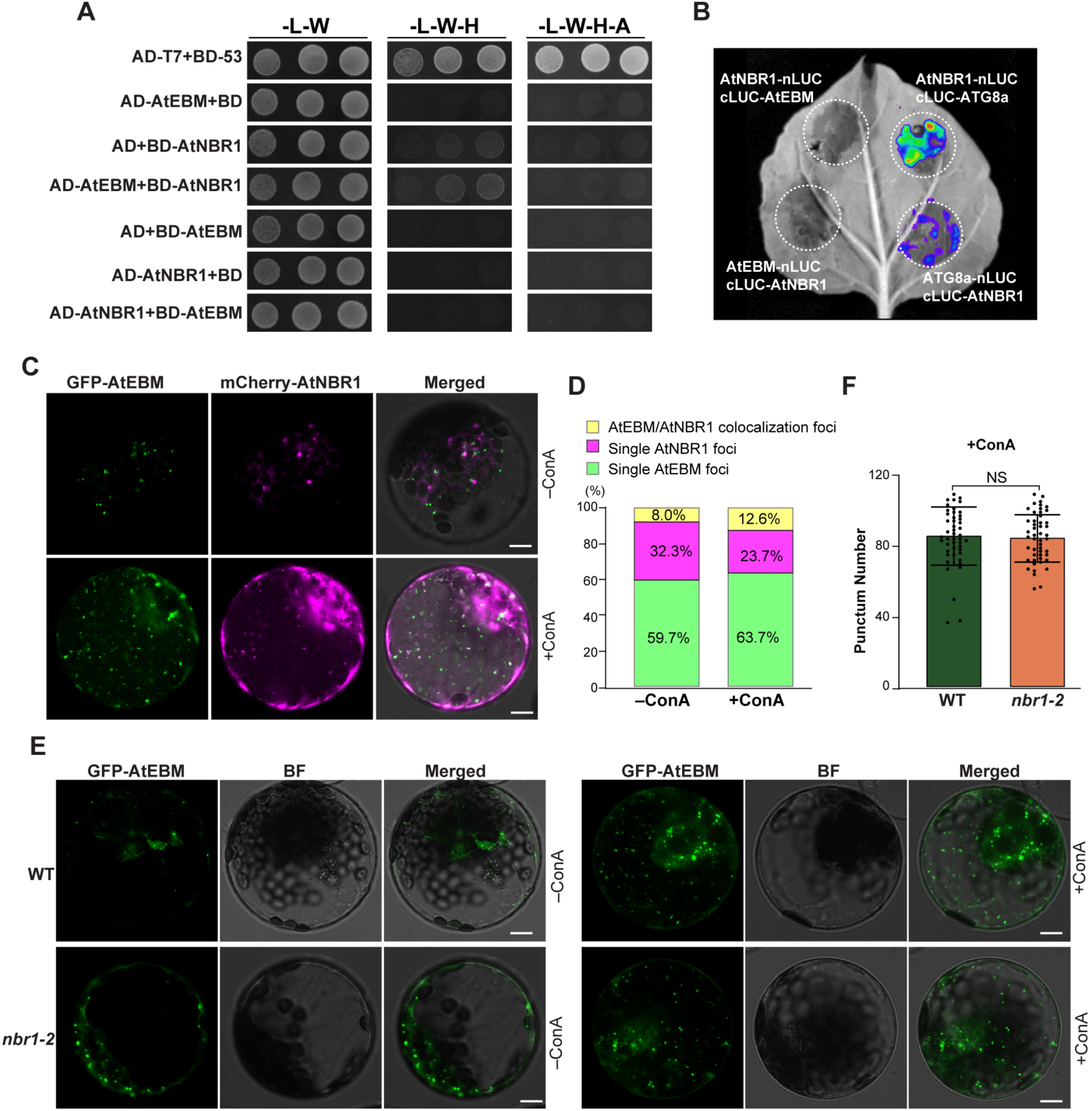
NBR1 is not essential for the vacuolar trafficking of AtEBM. **(A)** Yeast Two-Hybrid (Y2H) assay to test the interaction between AtEBM and AtNBR1. Full-length AtEBM and AtNBR1, expressed as N-terminal fusions to either the activation domain (AD) or the binding domain (BD), were coexpressed in yeast on a selection medium lacking leucine (Leu), tryptophan (Trp), histidine (His), and adenine (Ade) (-L-W-H-A), or on a nonselective medium lacking Leu and Trp (-L-W) as a control. AD and BD indicated empty AD and BD plasmids, respectively. **(B)** Luciferase complementation imaging (LCI) assay to test the interaction between AtEBM and AtNBR1. The CaMV 35S promoter-driven construct pairs indicated in the figure were infiltrated into different leaf regions of four-week-old *N. benthamiana* plants. The signal was detected 48 hours later. nLUC and cLUC indicated the N-terminal and C-terminal fragments of firefly luciferase, respectively. **(C)** Wild-type protoplasts were co-transformed with GFP-AtEBM and mCherry-AtNBR1, and then treated with either 1 μM ConA (+ConA) or DMSO (–ConA) for 12 to 14 hours before confocal imaging analysis. **(D)** Quantification of GFP-AtEBM puncta that co-localize with mCherry-AtNBR1. Images (*n* = 11-20) similar to those shown in (C) were analyzed to calculate co-localization ratios. **(E-F)** Leaf protoplasts from WT and *Arabidopsis nbr1-2* mutant were transformed with GFP-AtEBM and treated with 1 μM ConA (+ConA) or DMSO (–ConA) for 12 to 14 hours before confocal imaging analysis. Bars = 10 μm. Quantification of the numbers of puncta per cells after ConA treatment is shown in (F). The data represent the average values calculated from three independent experiments and *n* = 16 protoplasts were randomly selected from each experiment. Data were analyzed with GraphPad Prism 6 using one-way ANOVA followed by a Tukey’s multiple comparisons test. NS indicates not significant, and error bars represent SD.

Lastly, to further examine the subcellular localization of GFP-AtEBM, we coexpressed it with various markers of endocytic compartments, including the ER (CNX-mCherry), the Golgi stacks (Man1-RFP), the trans-Golgi network (RFP-SYP61), the MVB/PVC marker (mCherry-Rha1), the late endosome/vacuole (mCherry-RabG3f), and the tonoplast (mCherry-VAMP711). However, we failed to detect any overlapping signals between GFP-AtEBM and these markers (see Supplemental Figure 2), which suggests that the MVB trafficking pathway is unlikely to be involved in the vacuolar transport of AtEBM.

### AtEBM is an interacting partner of ATG8 mediated via the AIM-LDS interface

To further investigate how AtEBM is transported to the vacuole, we conducted a Y2H screen to test for the interaction between AtEBM and all identified ATG components from *Arabidopsis*. Consistent with previous proteomic studies showing that AtEBM was present in the immunoprecipitates of FP-tagged ATG8 (Zeng *et al*., 2021; Zhao *et al*., 2022; Zhou *et al*., 2023), we found that AtEBM could interact directly with AtATG8a by Y2H assay (Figure 4A), as was evident by the growth of yeast cells on the quadruple selection medium (SD-Ade-His-Trp-Leu). The interaction was further confirmed by using an LCI assay *in planta* and by an *in vitro* pull-down assay (Figure. 4B and 4C).

**Figure 4.**
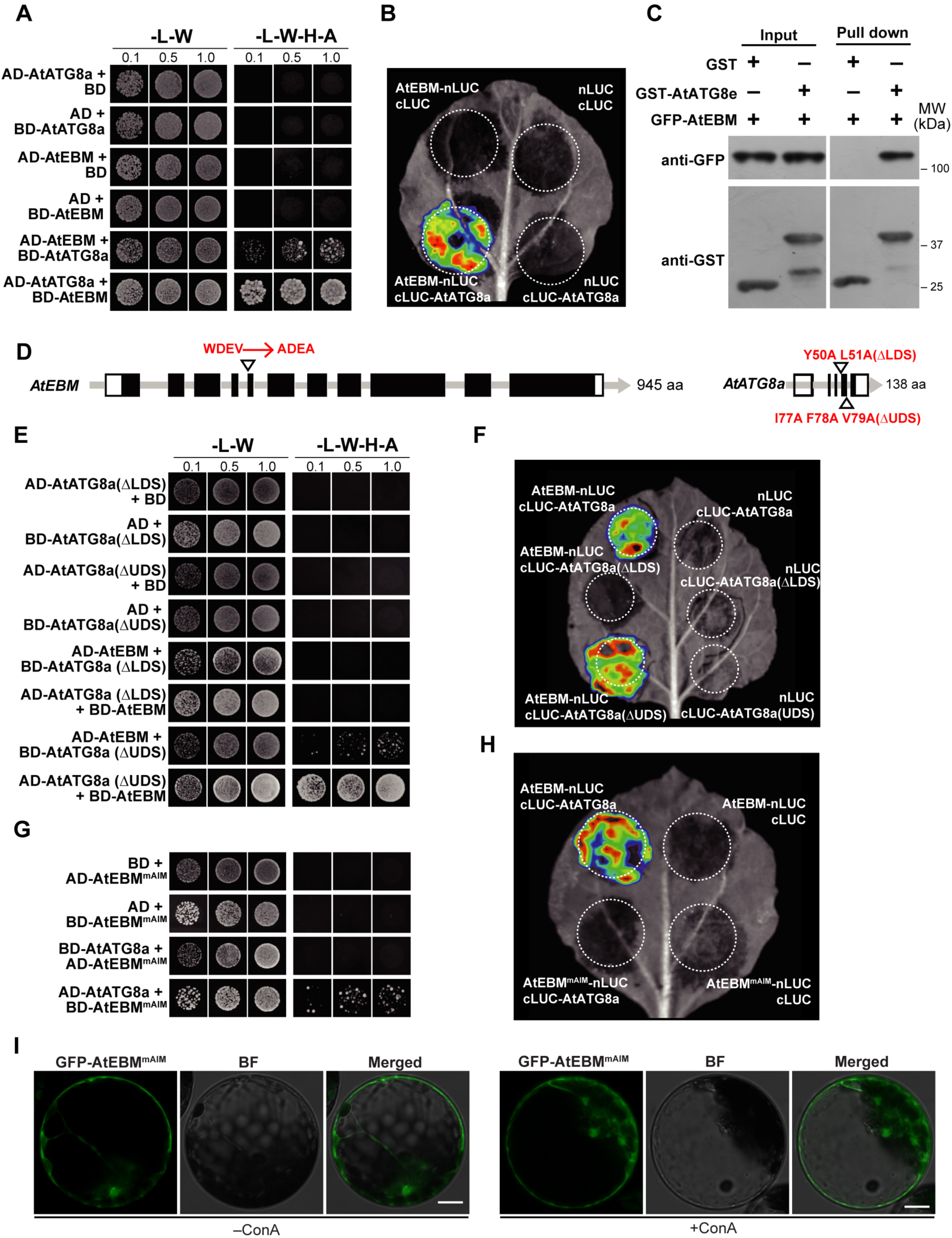
AtEBM interacts directly with AtATG8 via a conserved AIM motif. **(A-C)** Y2H, LCI, and co-immunoprecipitation (Co-IP) assays to test the interaction between AtEBM and AtATG8a. The Y2H and LCI assays were conducted as described in Figures 3A and 3B, respectively. For the Co-IP assay, plant lysates extracted from transgenic lines expressing GFP-AtEBM were used for IP with GST-AtATG8e or GST resins. The IP fractions were analyzed for the AtEBM-AtATG8 interaction by immunoblot analysis with the indicated antibodies. **(D)** Schematic diagram of the domain architectures of AtEBM and AtATG8a. AtEBM has a predicted AIM motif (W-D-E-V) at position 198 aa. The LDS and UDS are the LIR/AIM and UIM-docking sites of AtATG8a, respectively. **(E-F)** Y2H and LCI assays to reveal that AtEBM binds to AtATG8a in an LDS-dependent manner. Point mutations in the LDS, but not the UDS, of AtATG8a significantly reduced its binding to AtEBM. **(G-H)** Y2H and LCI assays to identify the AIM essential for AtEBM binding to AtATG8a. Point mutations of the predicted AIM of AtEBM at positions 198 and 201 by alanine substitutions (W198A and V201A) significantly reduced its binding to AtATG8a. **(I)** Subcellular localization of GFP-AtEBM^mAIM^ in *Arabidopsis* protoplasts. Leaf protoplasts from WT were transformed with GFP-AtEBM^mAIM^ and treated with 1 μM ConA (+ConA) or DMSO (–ConA) for 12 to 14 h before confocal imaging. Bars = 10 μm.

In general, ATG8-interacting proteins utilize two different motifs, the ATG8-interacting motif [AIM, also known as the LC3-interacting region (LIR)] or the ubiquitin-interacting motif (UIM), to bind to ATG8 at its LIR-docking site (LDS) or UIM-docking site (UDS), respectively (Marshall *et al*., 2019). To determine how AtEBM binds to ATG8, we designed two mutated variants of AtATG8a: one with alanine substitutions in the LDS [ATG8(ΔLDS), Y50A L51A] and the other with substitutions in the UDS [ATG8(ΔUDS), I77A F78A V79A] (Figure 4D). We then tested their interactions with AtEBM. As shown in Figures 4E and 4F, both Y2H and LCI assays showed that AtEBM could still bind to the ATG8(ΔUDS) variant but not the ATG8(ΔLDS) mutant.

We further performed an iLIR analysis (Kalvari et al., 2014; https://ilir.warwick.ac.uk) and identified a potential AIM [WDEV, amino acids (aa) 198-201] (Figure 4D). To test the essentiality of the predicted AIM for ATG8 binding, we mutated its key residues (AtEBM^mAIM^: W198A, V201A) to alanine within the full-length AtEBM protein. The Y2H and LCI results revealed that the AIM mutation completely abolished the interaction between AtEBM and AtATG8a (Figures 4G and 4H). Additionally, we examined the vacuolar trafficking of the GFP-AtEBM^mAIM^ fusion protein by transiently expressing it in *Arabidopsis* protoplast. Confocal imaging revealed that the AIM mutation significantly impaired AtEBM vacuolar trafficking, as no GFP-AtEBM-labelled puncta were detected in the vacuole, even after ConA treatment (Figure 4I). Taken together, these results indicated that AtEBM interact directly with ATG8a via an AIM-LDS interface, which is essential for its vacuolar trafficking.

### The conservation of AIM among EBM orthologs across the green lineage

Given the key role of AIM in the vacuolar trafficking of AtEBM, we explored the evolutionary origin of EBM proteins, as well as the functional conservation and diversification of the AIM (WDEV) among EBM orthologs. To this end, we reconstructed a phylogenetic tree of EBM orthologs alongside with multiple fungal and metazoan exo-type β-mannosidases, using a β-galactosidase (AT3G54440) as the outgroup (see Supplemental Figure 3A). Phylogenetic analysis suggests that the endo-type β-mannosidase first arose in green algae of the Chlorophyta phylum, such as *Tetraselmis striata* and *Chloropicon primus*. It then spread to most green plants including unicellular chlorophytes, streptophytes, bryophytes, lycophytes, gymnosperms, and angiosperms (see Supplemental Figure 3B). Interestingly, some chlorophyte algae (*e.g.*, *T*. *striata* and *Coccomyxa subellipsoidea*) and streptophytes (*e.g.*, *Chara braunii* and *Klebsormidium nitens*) appear to have both endo- and exo-type β-mannosidases, whereas most green plants only have the endo-type (Figure 5A and Table S3). Using the iLIR program to analyze these putative EBM orthologs, we found that the WDEV motif is present in all EBM orthologs of land plants, though it tends to be less perfectly conserved in early plant species. Conversely, the corresponding regions of exo-type β-mannosidases are much more diverse than those of endo-type enzymes (Figure 5A).

**Figure 5.**
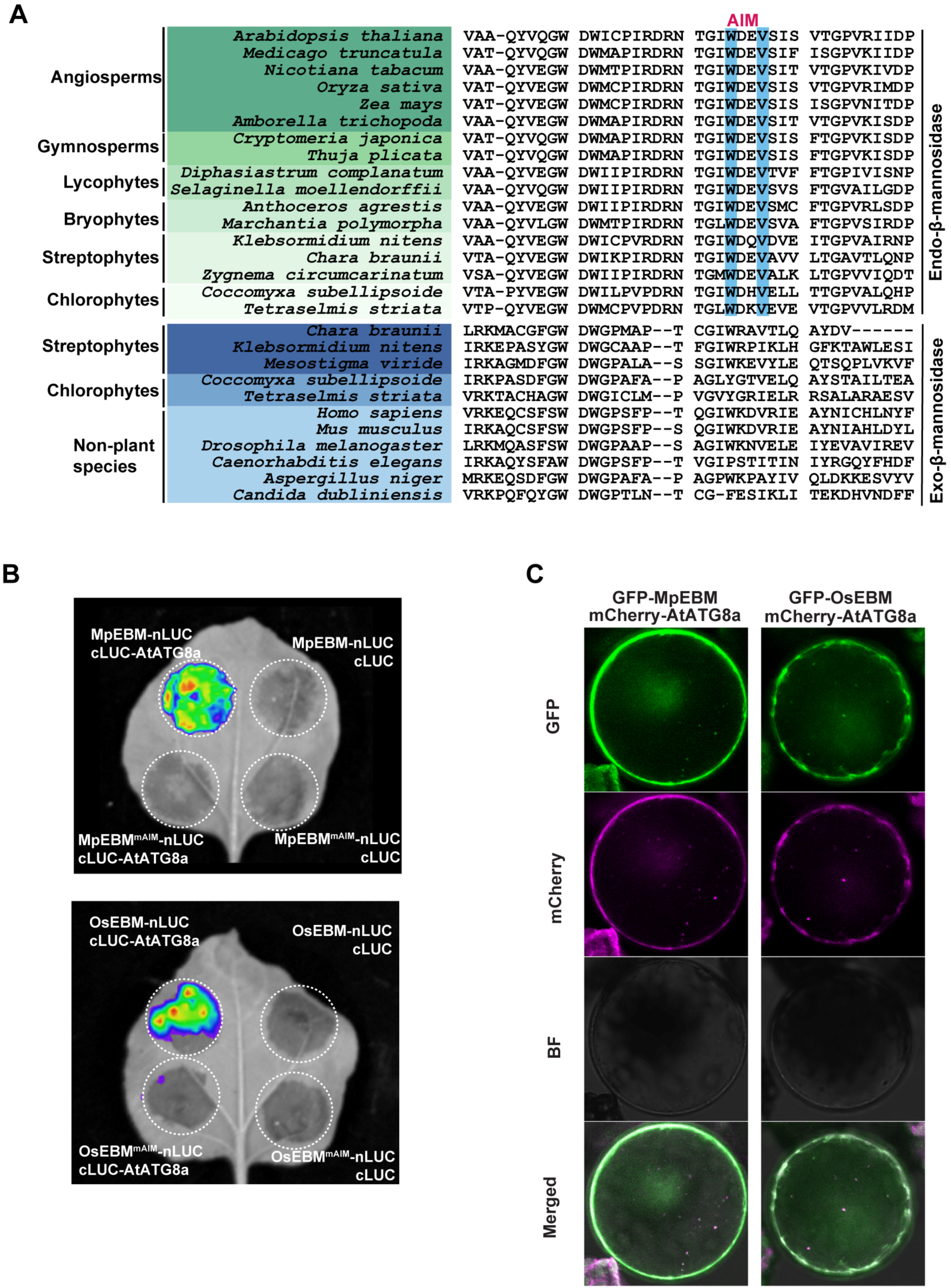
Functional conservation and diversity of the AIM sequences in EBM orthologs across the green lineage. **(A)** Sequence alignment of the predicted AIM regions of the EBM orthologs from representative green lineage species and the corresponding regions of exo-type β-mannosidases from early plants and non-plant species. The key conserved tryptophan (W) residue and valine (V) residues in the putative AIMs are highlighted in blue. **(B)** LCI assay to test the interaction between EBM and ATG8 orthologs from liverwort (MpEBM) and rice (OsEBM). The assay was conducted as described in Figure 3B. **(C)** Protoplast transient expression assay to examine the subcellular localization of EBM orthologs from liverwort and rice. The assay was conducted as described in Figure 1.

To determine the extent to which the EBM can interact with ATG8 orthologs across the green lineage, we selected EBM orthologs from rice and liverwort. We then tested their interactions with ATG8 using an LCI assay. As shown in Figure 5B, both EBM proteins interacted strongly with their corresponding ATG8 orthologs. However, their AIM-mutated variants could not interact. A protoplast transient expression assay demonstrated that, similar to AtEBM, MpEBM from *M. polymorpha* and OsEBM from *O. sativa* also colocalized well with the autophagy marker ATG8 after ConA treatment (Figure 5C). Taken together, these results provide the evidence for the evolutionary conservation of the interaction between EBMs and ATG8 and of autophagy-mediated trafficking of EBMs.

### Autophagy mediates the vacuolar trafficking of GFP-AtEBM

To further illustrate the vacuolar trafficking pathway of AtEBM, we generated transgenic *Arabidopsis* plants that express GFP-AtEBM under the control of its native promoter as well as the *Arabidopsis UBIQUITIN 10* (*UBQ10*) gene promoter. We then analyzed their distribution patterns in root cells. After subjecting the seedlings to N starvation and ConA treatment, we readily detected numerous GFP-AtEBM-labelled puncta in the vacuoles of *proAtEBM::GFP-AtEBM* transgenic lines, albeit the signal intensity was relatively low (see Supplemental Figure 4). This finding is consistent with the observations from protoplast transient expression assays (Figures 1 and Figure 2). The subcellular distribution pattern of the *proUBQ10::GFP-AtEBM* reporter in transgenic lines was similar to the *proAtEBM::GFP-AtEBM* reporter (Figure 6A). When *proUBQ10::GFP-AtEBM* reporter was introgressed into the autophagy-deficient mutant *atg7-2*, the accumulation of GFP-AtEBM-decorated puncta in the vacuole was completely blocked (Figure 6A). Using colocalization studies, we confirmed that GFP-AtEBM-labelled puncta were autophagic vesicles by coexpressing GFP-AtEBM and the autophagic marker mCherry-ATG8e (Zhuang *et al*., 2013). Figure 6B and 6C show that, of all the imaged GFP-AtEBM puncta, 56% and 73%, respectively, were decorated with mCherry-ATG8e before and after ConA treatment. To further confirm these observations, we examined the vacuolar cleavage of GFP-AtEBM in wild-type and *atg7-2* seedlings. When GFP-tagged fusion proteins are targeted to the vacuole for degradation, the free GFP moiety is released and can be detected by immunoblotting analysis with anti-GFP antibodies. In wild-type seedlings, the free GFP band was easily detected whereas the full-length GFP-AtEBM fusion protein was barely detectable. Its intensity increased with ConA treatment. In contrast, only GFP-AtEBM fusion protein was found in the *atg7-2* mutant (Figure 6D). This pattern is similar to that of the GFP-ATG8A described previously (Suttangkakul *et al*., 2011), suggesting that autophagy mediates the vacuolar trafficking of the GFP-AtEBM fusion protein.

**Figure 6.**
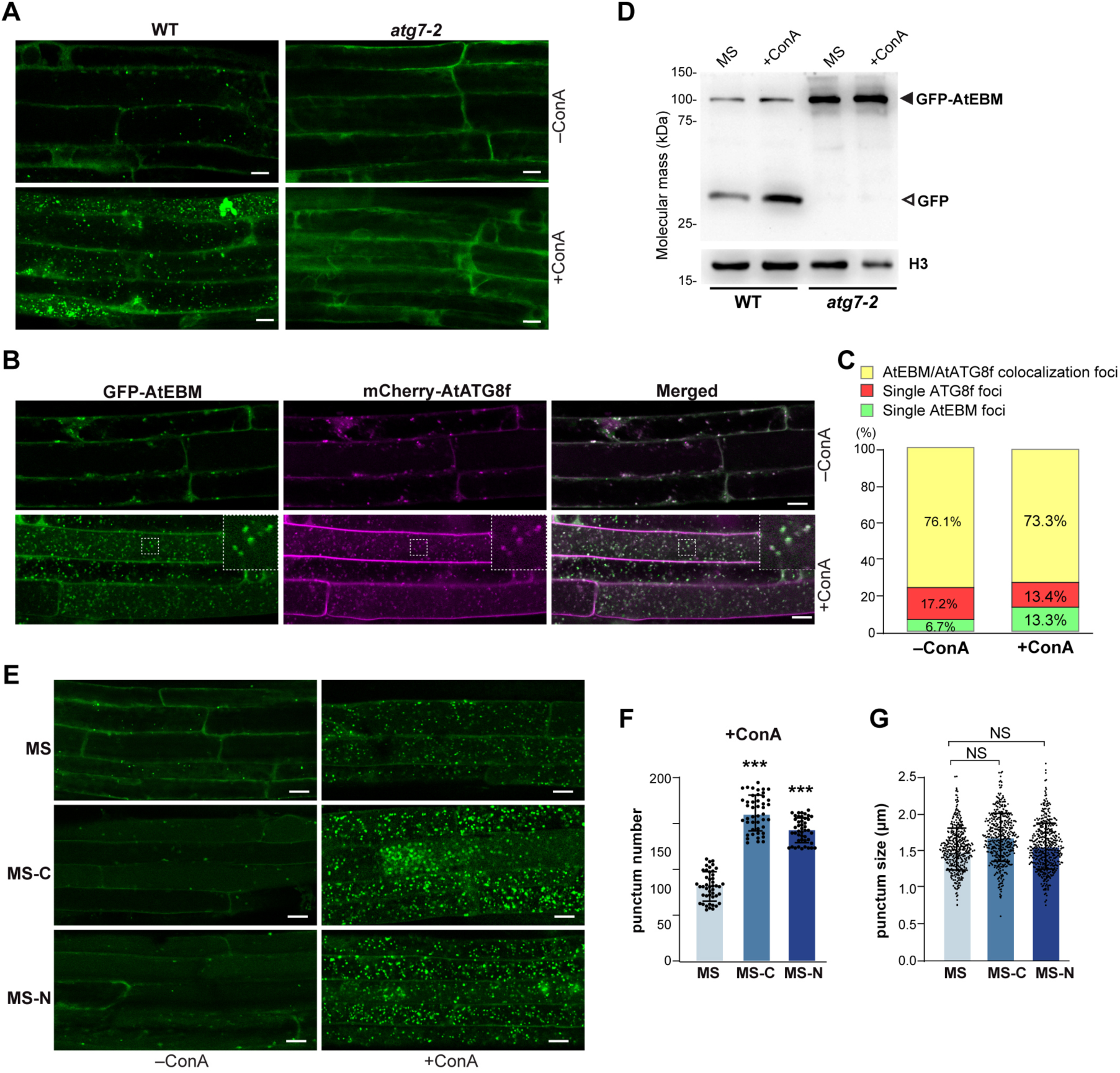
GFP-AtEBM is delivered to the vacuole via the autophagic pathway. **(A)** Confocal imaging of the root cells expressing GFP-AtEBM. The GFP-AtEBM with the indicated genetic backgrounds were grown on MS medium for 6 days and then transferred to nitrogen-free medium containing 1 μM ConA or DMSO (–ConA) for 12 hours before confocal microscopy. Bars = 10 μm. **(B)** GFP-AtEBM colocalizes with mCherry-AtATG8f in autophagic vesicles. Wild-type plants stably expressing both reporters were grown on MS medium for 6 days and then transferred to fresh MS medium containing 1 μM ConA or DMSO (–ConA) for 12 hours before confocal microscopy. Insets show 3x magnifications of the dashed boxes. Bars = 10 μm. **(C)** Quantification of the colocalization ratio between GFP-AtEBM and mCherry-ATG8f. In total, *n* = 313 (–ConA) and *n* = 939 (+ConA) particles were used for colocalization ratio calculations. **(D)** GFP-AtEBM cleavage assay. WT or *atg7-2* seedlings expressing GFP-AtEBM were treated as described above, and total protein was extracted for immunoblot analysis using anti-GFP bodies. Antibody against histone H3 was used as a loading control. **(E)** Deposition of autophagic bodies containing GFP-AtEBM inside the vacuole under nutrient-depleted conditions. Seedlings expressing GFP-AtEBM were grown for 6 days on MS solid medium with 1% Suc (w/v) and then transferred to fresh MS, sucrose- (–C) or nitrogen-deficient (–N) liquid media with or without the addition of 1 μM ConA, for 8 hours before confocal observation. Bars = 10 μm. **(F)** Quantification of the numbers of puncta per section in the root cells in (E). The data represent the average values calculated from three independent experiments; *n* = 45 sections per treatment. **(G)** Quantification of the size of the GFP-AtEBM-positive autophagic bodies in (E). The data represent the mean values calculated from three independent experiments. *n* > 500 autophagic bodies per treatment were randomly selected from at least 15 different regions within five sections from each of three independent experiments. Data in (F) and (G) were analyzed using GraphPad Prism 6 with one-way ANOVA followed by Tukey’s multiple comparisons test. NS indicates not significant. *** indicates *p* < 0.01. Error bars represent SD.

It is known that the yeast ScApe1 is transported to the vacuole by Cvt vesicles under growing conditions in nutrient-rich medium, and by the autophagosomes during starvation (Baba *et al*., 1997). Since AtEBM transport occurs constitutively under growing conditions (Figure 6A), we wondered whether its transport is affected by nutritional status. Therefore, we examined the seedlings expressing GFP-AtEBM after transferring them to nutrient-limiting media. Figure 6F shows that, unlike under normal growing conditions, a ∼2-fold increase in the number of GFP-AtEBM-labeled puncta was detected in starved root cells. Under MS conditions, the size of GFP-AtEBM-labeled puncta is fairly homogeneous, calculated to be 1.3 μm in diameter. This size is similar to that of puncta under starved conditions (Figure 6G) and to the size of previously reported GFP-ATG8-labeled puncta (Huang *et al*., 2019). These observations strongly suggests that AtEBM is likely transported to the vacuoles via autophagosomes regardless of nutrient availability.

### Phenotypic analyses demonstrate that AtEBM is not essential for bulk autophagy but required for plant fertility

To uncover the function of *AtEBM*, we first analyzed its expression profiles based on the information from publicly available RNA-seq databases, including TraVA (Transcriptome Variation Analysis, http://travadb.org/) and ARS (Arabidopsis RNA-seq Database, http://ipf.sustech.edu.cn/pub/athrna/). As shown in Supplemental Figure 5A, *AtEBM* was expressed ubiquitously in most tissues across different developmental stages, with slightly higher expression in mature or senescent organs. This was further validated by analyzing GUS activities in *proAtEBM::GUS* transgenic seedlings (see Supplemental Figure 5B).

Additionally, we constructed a coexpression network of *AtEBM* using ATTED-II version12 (http://atted.jp/top_draw.shtml). Consistent with our previous observations, the coexpression analysis revealed that *AtEBM* is tightly coexpressed with several key components of the autophagy pathway, including *ATG8f*, *ATG13a*, and *VPS15* (see Supplemental Figure 5C). Furthermore, *AtEBM* was coexpressed with multiple genes involved in glycan degradation and proteolysis, suggesting its potential involvement in *N*-glycan hydrolysis and protein degradation processes.

Next, we used CRISPR/Cas9-mediated genome editing with guide RNAs targeting the first and second exons of *AtEBM* to generate mutants (see Supplemental Figure 6>A). This mutagenesis generated two independent *AtEBM* alleles, designated *atebm-1* and *atebm-2*, which harbor either 360- and 287-bp deletions, respectively (see Supplemental Figures 6B and 6C). The two deletion mutations introduced reading frameshifts downstream of the Thr-19 and Val-46 codons, respectively, which should substantially truncate the AtEBM protein if translated (see Supplemental Figure 6D).

Given that AtEBM coexpresses well with several key ATG components (Supplemental Figure 5C), we first examined the growth phenotypes of the *atebm* mutants and challenged them with multiple nutrient-limiting conditions to assess the role of *AtEBM* in autophagy. When grown under nutrient-rich conditions, the *atebm* homozygous mutants resembled the wild type and did not exhibit early leaf senescence, a typical phenotype of autophagy mutants, such as the *atg5-1* and *atg7-2* mutants (see Supplemental Figure 7). Furthermore, as shown in Supplemental Figures 8 and 9, the *atebm* mutants responded nearly indistinguishably from wild-type plants under all tested nutrient stress conditions, and were insensitive to either nitrogen starvation or fixed-carbon limitation.

In addition to examining the phenotypes of *atebm* mutants under nutrient stress conditions, we also examined them under several other abiotic stress conditions, such as salinity, osmotic, and heat stress. Autophagy mutants (*atg5-1* and *atg7-2*) exhibited reduced responses to these stresses, while no significant differences were observed between wild type and *atebm* mutants (data not shown). This finding is similar to a previous study on the *Arabidopsis atfuc1* mutant, which is defective in α1,3/4-fucosidase activity that is essential for *N*-glycan degradation (Kato *et al*., 2018). Taken together, it is likely that AtEBM is an autophagy cargo rather than a component involved in bulk autophagy.

During the phenotyping process, we noticed that *atebm* mutants displayed certain abnormalities in silique length and seed set (Figure 7). 45-day-old plants grown at 22°C under LD conditions were scored for the silique length on the main inflorescence. Siliques were then divided into three major categories based on fertility: fully fertile (Type I), partially sterile (Type II), and completely sterile (Type III). Under favorable growth conditions, wild-type plants only produced Type II (1.46%) and Type III (1.01%) siliques marginally. Conversely, *atebm* plants produced much higher percentages of Type II and Type III siliques (8.76% Type II and 6.39% Type III siliques in *atebm-1* and 10.11% Type II and 8.99% Type III siliques in *atebm-2*, respectively) (Figures 7B and 7C). Such silique defects in *atebm* mutants are not due to pollen viability since *in vitro* and *in vivo* pollen germination assays showing that the *atebm* mutants could produce highly viable pollens similar to *atg* mutants (Supplemental Figure 10). Interestingly, the frequency of silique defects in *atebm* mutants was also higher than those of *atg5-1* and *atg7-2* mutants, which further implying that such growth defects are not associated with the autophagy pathway and are likely related to the other functions of AtEBM.

**Figure 7.**
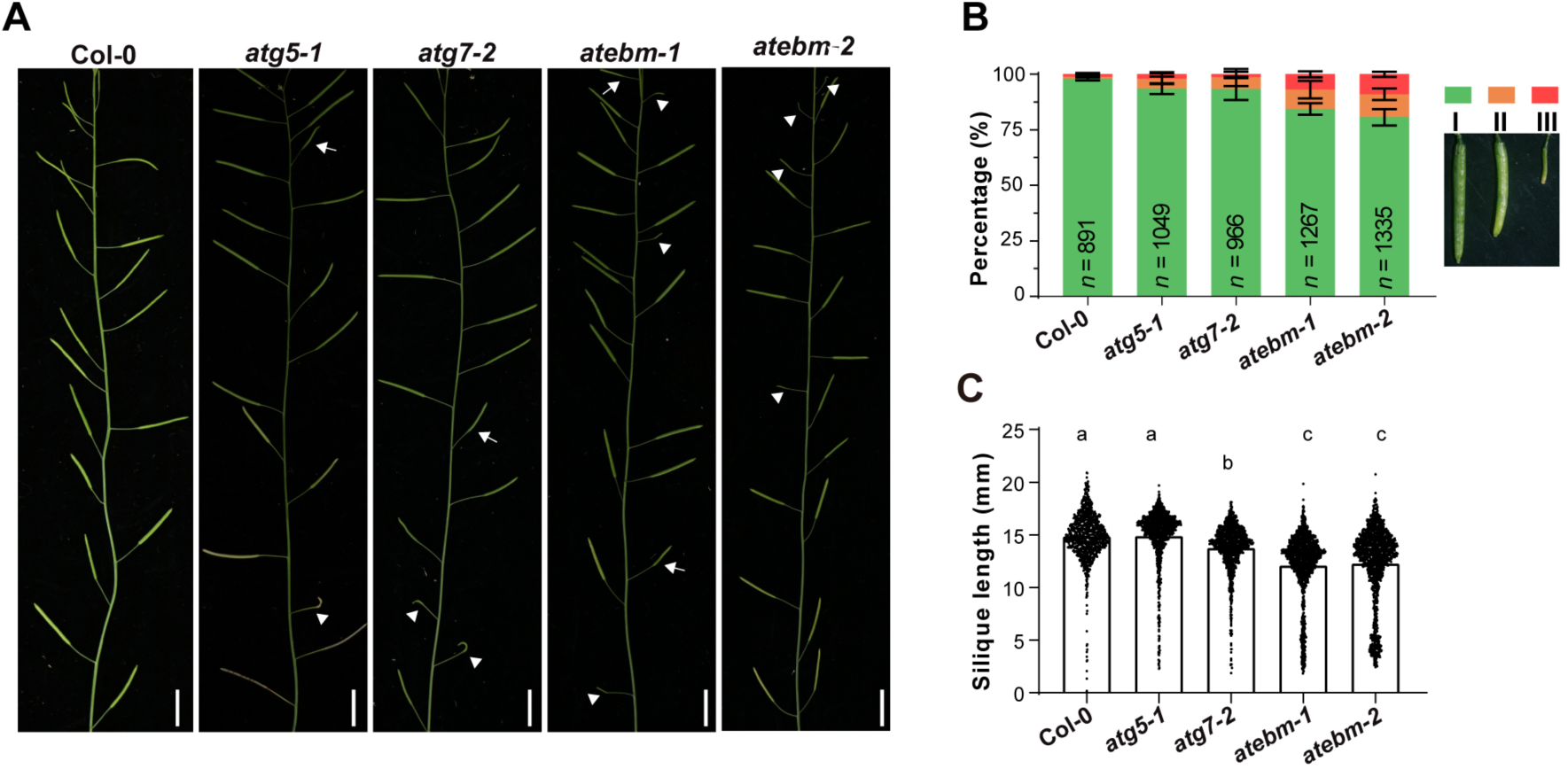
Silique phenotypes of *atebm* mutants. **(A)** Typical siliques of the primary inflorescence in WT, *atg* mutants, and *atebm* mutants grown at 22°C for 45 days under LD conditions. Siliques were divided into three categories: fully fertile (Type I), partially sterile (Type II), and completely sterile (Type III). The white arrows indicate Type II siliques and the white arrowheads indicate Type III siliques, respectively. **(B)** The percentages of each type of silique in WT, *atg* mutants, and *atebm* mutants. **(C)** Violin plot shows the silique length of WT, *atg* mutants, and *atebm* mutants. The data represent the average values calculated from three independent experiments; *n* > 800 per genotype.

## DISCUSSION

The vacuole, the major catabolic organelle in budding yeast and plants, is essential for cell homeostasis, nutrient storage and recycling, and detoxification. It harbors many evolutionarily conserved transporters and hydrolases dedicated to these functions (Hecht *et al*., 2014; Shimada *et al*., 2018). Three trafficking routes to this organelle have been well characterized in budding yeast: the CPY pathway, the AP-3 pathway, and the Cvt pathway (Hecht *et al*., 2014; Eising *et al*., 2022). Similar protein sorting pathways to the CPY and AP-3 pathways have been identified in plants and are being characterized (Di Sansebastiano *et al*., 2018; Aniento *et al*., 2022). In this study, we found that the vacuolar glycosidase AtEBM in *Arabidopsis* is transported from the cytoplasm to the vacuole via the autophagy pathway. Since the glycosidase Ams1 in yeast is a cargo of the Cvt pathway, our finding strengthens the similarity and the evolutionary relatedness of the vacuolar protein targeting pathways in budding yeast and plants. Given that *Arabidopsis* and budding yeast are two distant species that diverged more than over billion years ago (Wang *et al*., 1999), it is likely that autophagy in other kingdoms or phyla might also serve as a route for the vacuolar trafficking of hydrolase cargoes.

*S. cerevisiae* ScAtg19/ScAtg34 and *S. pombe* SpNbr1 were employed as autophagy receptors that mediate the vacuolar trafficking of hydrolase cargoes in the Cvt and the NVT pathways, respectively (Liu *et al*., 2015; Yamasaki & Noda, 2017). Although the SpNbr1 ortholog is present in the *Arabidopsis* genome (Svenning *et al*., 2011), the AtEBM transport seems to be independent of AtNBR1, as evident from the lack of interaction between AtEBM and AtNBR1, the minimal colocalization between GFP-AtEBM- and mCherry-AtNBR1-labelled puncta, and no alteration in accumulation of GFP-AtEBM puncta in the vacuoles of *nbr1-2* mutant compared to those of wild type (Figure 3). Furthermore, colocalization analysis with various markers of endocytic compartments revealed that, unlike in the NVT pathway, the vacuolar deposition of AtEBM was not associated with MVB sorting machinery (see Supplemental Figure 2). Instead, we found that AtEBM could directly interact with AtATG8 via an evolutionarily well-conserved AIM-LDS interface, which was essential for the vacuolar trafficking of EBMs (Figure 4). The fact that yeasts and plants use completely different approaches to recruit hydrolase cargoes suggests that the function of autophagy in the trafficking of vacuolar hydrolases can be maintained during evolution despite the loss of conservation in cargo recruitment.

Interestingly, some exo-type β-mannosidases, such as *Homo sapiens* Hs_MANBA and *Klebsormidium nitens* kfl00346_0110_v1.1, contain a less conserved WxxV motif in the corresponding regions (Figure 5A). Additionally, some lack typical signal peptides (see Supplemental Table S3). Therefore, it would be interesting to determine whether the autophagy machinery is involved in the vacuolar trafficking of these exo-type β-mannosidases.

In the Cvt pathway of *S. cerevisiae* and the NVT pathway in *S. pombe*, multiple cargoes commonly form a complex with autophagy receptors and are delivered together to the vacuole. For instance, ScApeI and ScAms1 bind to the receptor ScAtg19 and further assemble into the Cvt complex. Meanwhile, the soluble hydrolases SpLap2 and SpApe2 interact directly with SpNbr1 to form the NVT complex (Liu *et al*., 2015; Yamasaki & Noda, 2017). Therefore, it is tempting to speculate that AtEBM forms complexes with other cargoes and then are deposited to the vacuole together. Previous proteomic studies in *Arabidopsis* have identified over hundreds of hydrolases in vacuolar sap. Several of these hydrolases lack a typical signal peptide, including AtDAP1(AT5G60160, the homolog of yeast ScApe1), AtLAP1 (At2g24200) and AtLAP2 (At4g30920, the homologs of yeast ScLap3) (Carter *et al*., 2004; Jaquinod *et al*., 2007; Ohnishi *et al*., 2018). More work is needed to reveal how these hydrolases are transported to the vacuole and whether autophagy is involved in their vacuolar deposition.

Studies have shown that autophagy is also involved in the vacuolar transport of vacuolar processing enzymes (VPEs) in potato (*Solanum tuberosum*) leaves and in tobacco (*N. tabacum*) BY-2 suspension-cultured cells (Teper-Bamnolker *et al*., 2021). VPEs are cysteine-type endopeptidases that are synthesized as large precursor proteins in the ER. During carbon starvation, they are relocalized from the ER to autophagosomes and then transported to the vacuole via autophagy. However, the detailed mechanism by which VPEs are delivered to the vacuole via autophagy remains unclear, although several conserved potential AIMs have been predicted in VPEs (Teper-Bamnolker *et al*., 2021). Unlike the ER to vacuole trafficking of VPEs, AtEBM did not colocalize with the ER marker (Supplemental Figure 2), but rather overlapped well with the autophagy marker ATG8 in the cytosol (Figure 6B), suggesting that the AtEBM transport is more similar to the yeast Cvt pathway, which transports cargoes directly from the cytoplasm to the vacuole. In addition, like the Cvt pathway, AtEBM transport is a constitutive biosynthetic event that operates under nutrient-rich conditions, and the transport could be enhanced by nutrient stress (Figure6 E-G). However, the sizes of AtEBM-containing puncta were similar under nutrient-rich and -depleted conditions. This is different from the yeast ScApe1, which is transported to the vacuole by small Cvt vesicles under growing conditions and by large autophagosomes during starvation. In summary, this is the first report of a Cvt-like pathway employed in *Arabidopsis* to deliver the glycosidase AtEBM to the vacuole.

## ACKNOWLEDGEMENTS

This work was supported by grants from the National Natural Science Foundation of China (grant 32370352 to F.Q.L. and grant 32070195 to X.H.), the Natural Science Foundation of Guangdong Province (2024A1515011671) to X.H. We would like to thank Prof. Caiji Gao (South China Normal University, China) and Prof. Liwen Jiang (The Chinese University of Hong Kong, China) for the endocytic markers, and Prof. Qijun Chen (China Agricultural University, China) for the CRISPR/Cas9 reagents.

## AUTHOR CONTRIBUTIONS

F.Q.L. and L.N. designed the research. H.W. and J.Y.M. performed most of the experiments. J.Y.S., G.N.F., S.T.F., C.M.T., and Y.J.X. provided technical supports. F.Q.L., L.N., X.H., and H.W. wrote the manuscript with input from all authors.

## Data availability

The authors declare that all data supporting the findings of this study are available within the paper and its supplementary information, or are available from the corresponding author upon reasonable request.

## Supplemental Data

The following materials are available in the online version of this article.

**Supplemental Table 1.** Oligonucleotide Primers Used in the Study.

**Supplemental Table 2.** Vacuolar Glycosidases Identified in the Previous Proteomic Studies.

**Supplemental Table 3.** Protein Sequences of EBMs in the Plant Kingdom.

**Supplemental Table 4.** Coexpressed Genes Associated with *AtEBM*.

**Supplemental Figure 1.**
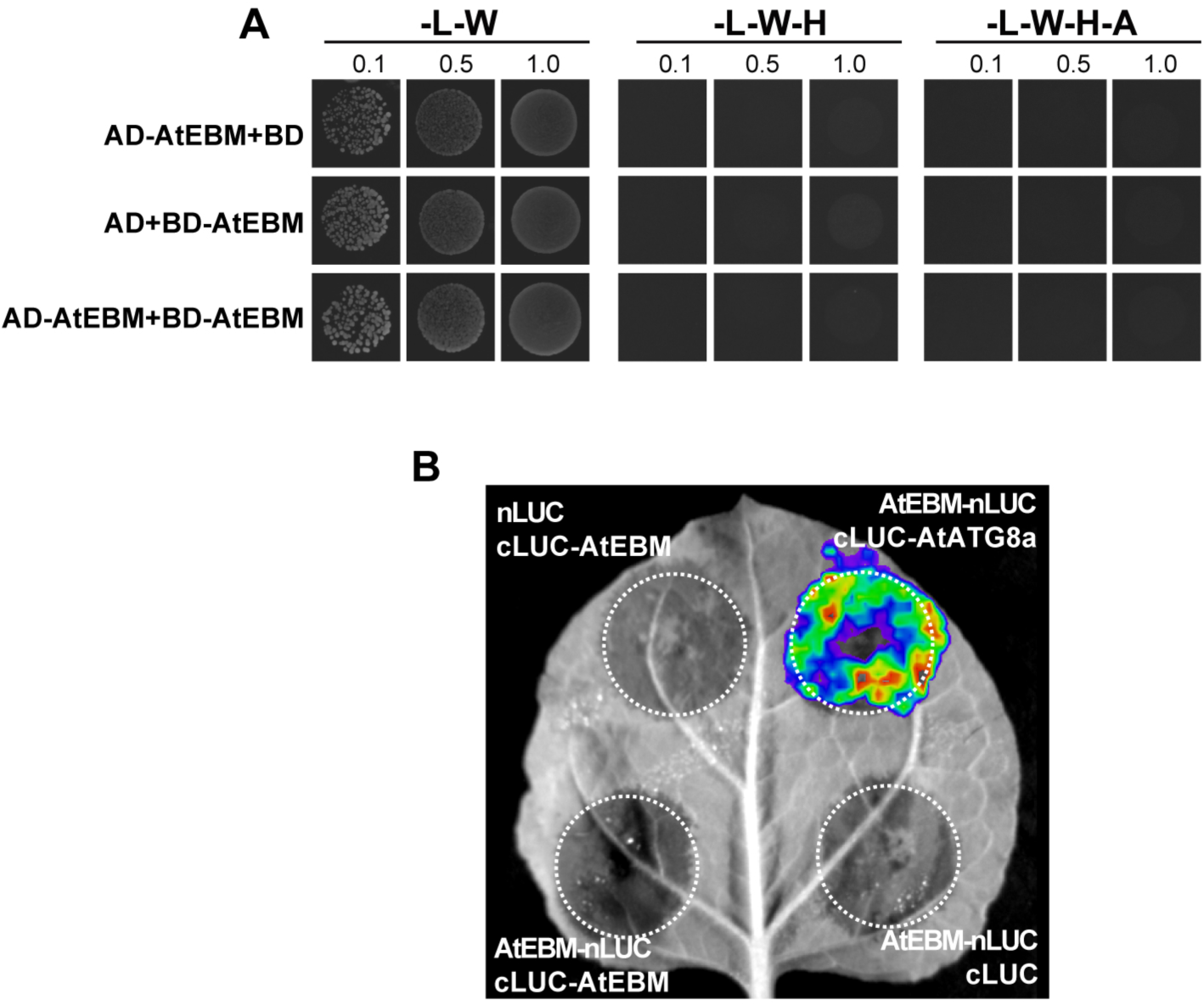
No self-interaction was observed for AtEBM. Y2H (A) and LCI assays (B) were performed as shown in Figure 3 to test the self-interaction of AtEBM. The interaction between AtEBM and AtATG8a was served as a positive control.

**Supplemental Figure 2.**
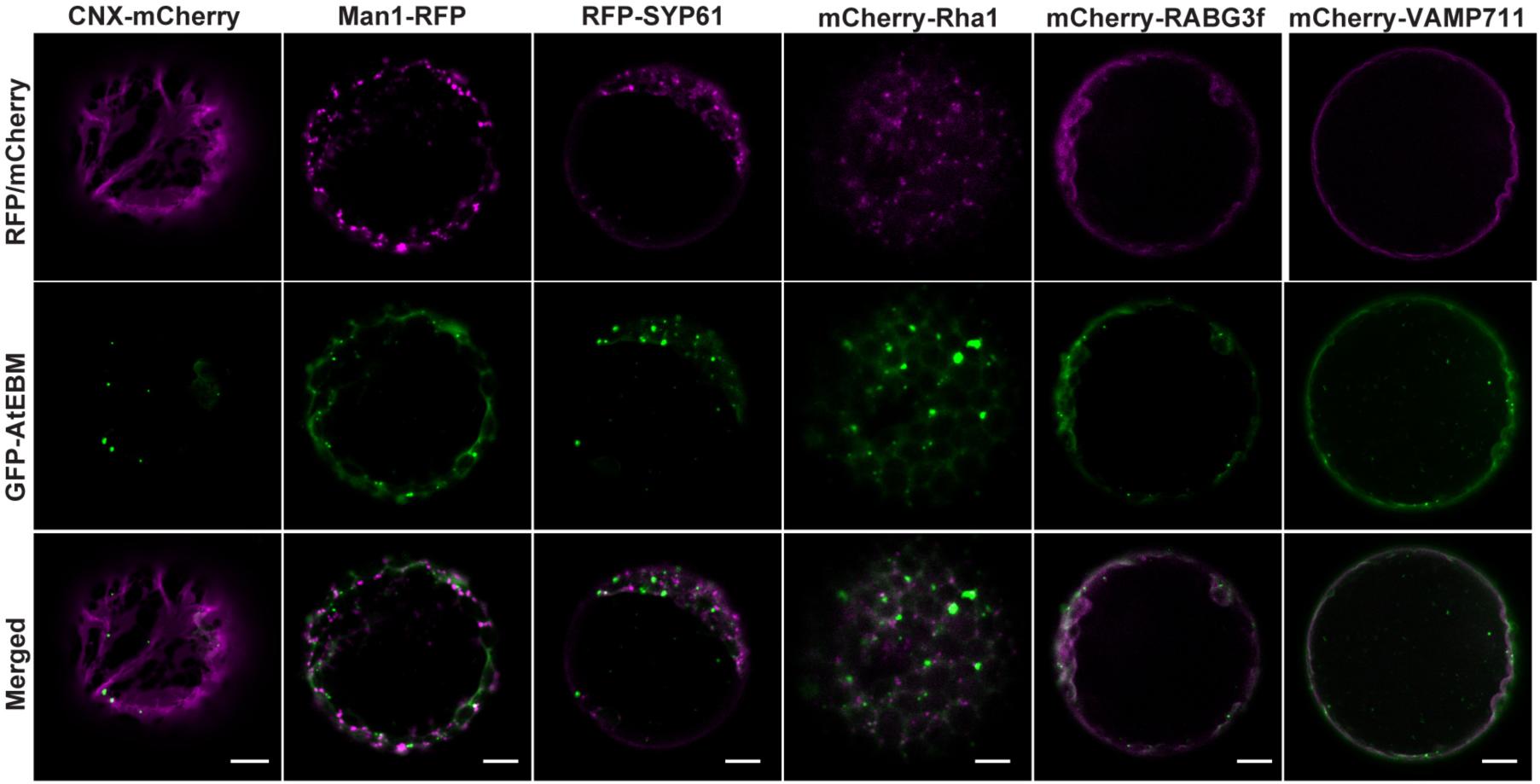
Colocalization analysis of AtEBM with various markers of endocytic compartments. GFP-AtEBM was coexpressed with a series of fluorescent organelle markers, followed by confocal microscopy at 12 to 14 hours after transfection. CNX-mCherry, ER marker; Man1-RFP, Golgi marker; RFP-SYP61, TGN marker; mCherry-Rha1, MVB/PVC marker; mCherry-RabG3f, late endosome/vacuole marker; mCherry-VAMP711, vacuolar membrane marker. Bars = 10 μm.

**Supplemental Figure 3.**
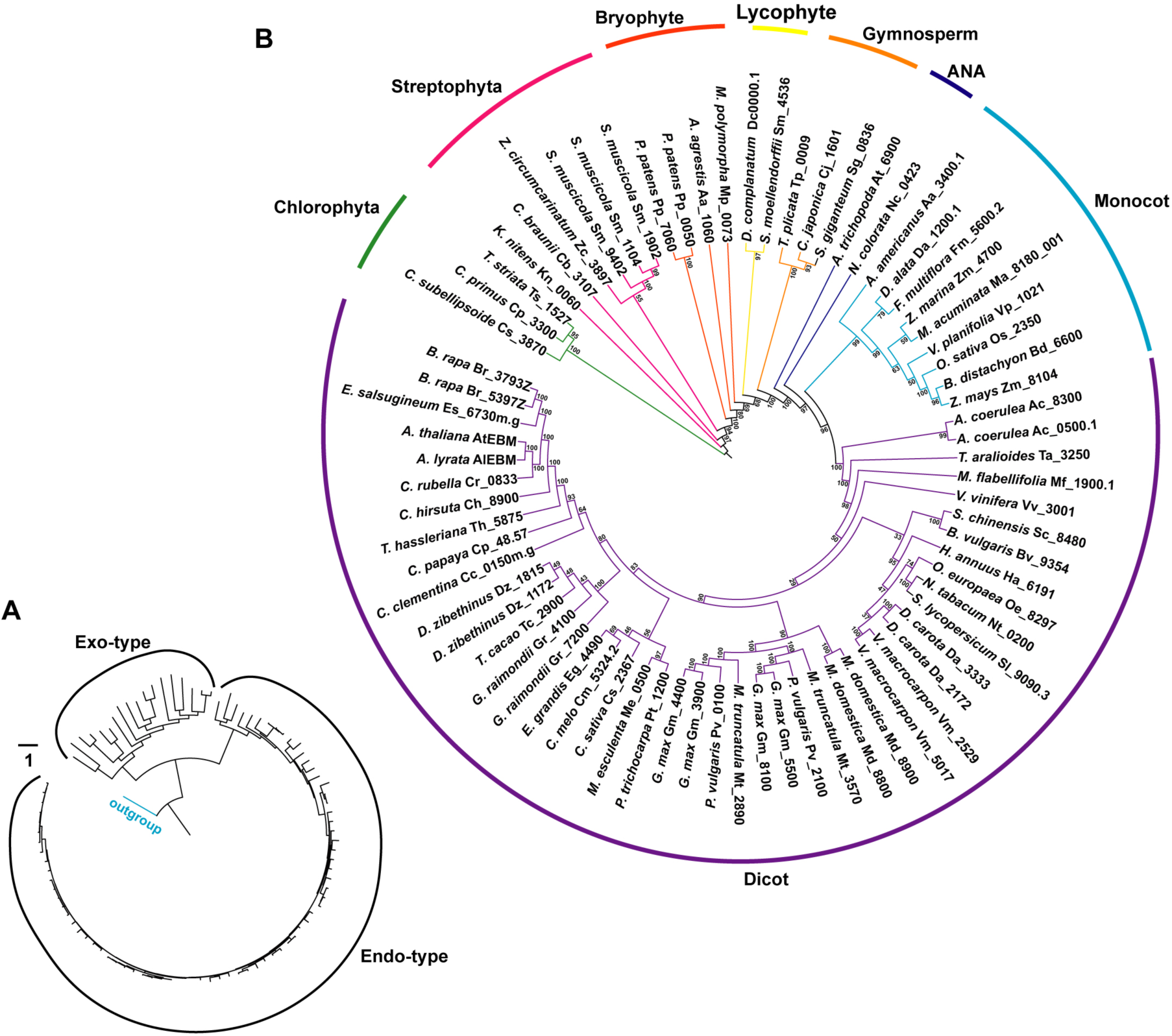
Genome-wide identification and phylogenetic investigation of EBMs in the plant kingdom. **(A)** Maximum likelihood tree topology for the endo- and exo-type β-mannosidases. *Arabidopsis* β-galactosidase (At3g54440) of the GH2 family was used as the outgroup. **(B)** Maximum likelihood tree topology for the endo-type β-mannosidases.

**Supplemental Figure 4.**
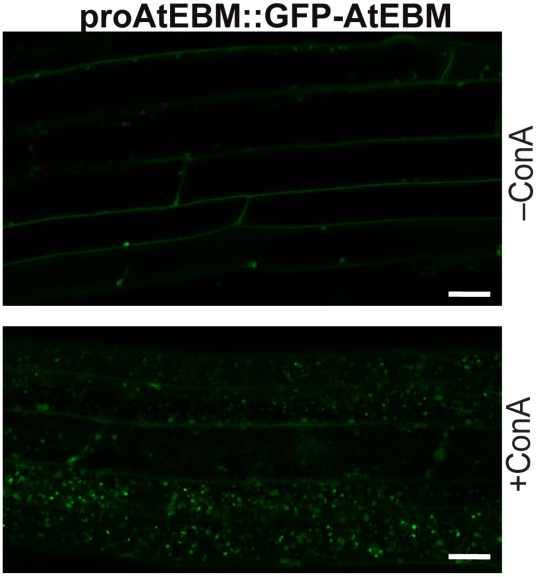
Subcellular locations of GFP-AtEBM in the root cells of transgenic seedlings expressing *proAtEBM::GFP-AtEBM*. Seeds of the *proAtEBM::GFP-AtEBM* transgenic lines were germinated and grown on MS medium for 6 days and then transferred to fresh medium containing 1 μM ConA (+ConA) or DMSO (–ConA) for 12 hours before confocal microscopy. Bars = 10 μm.

**Supplemental Figure 5.**
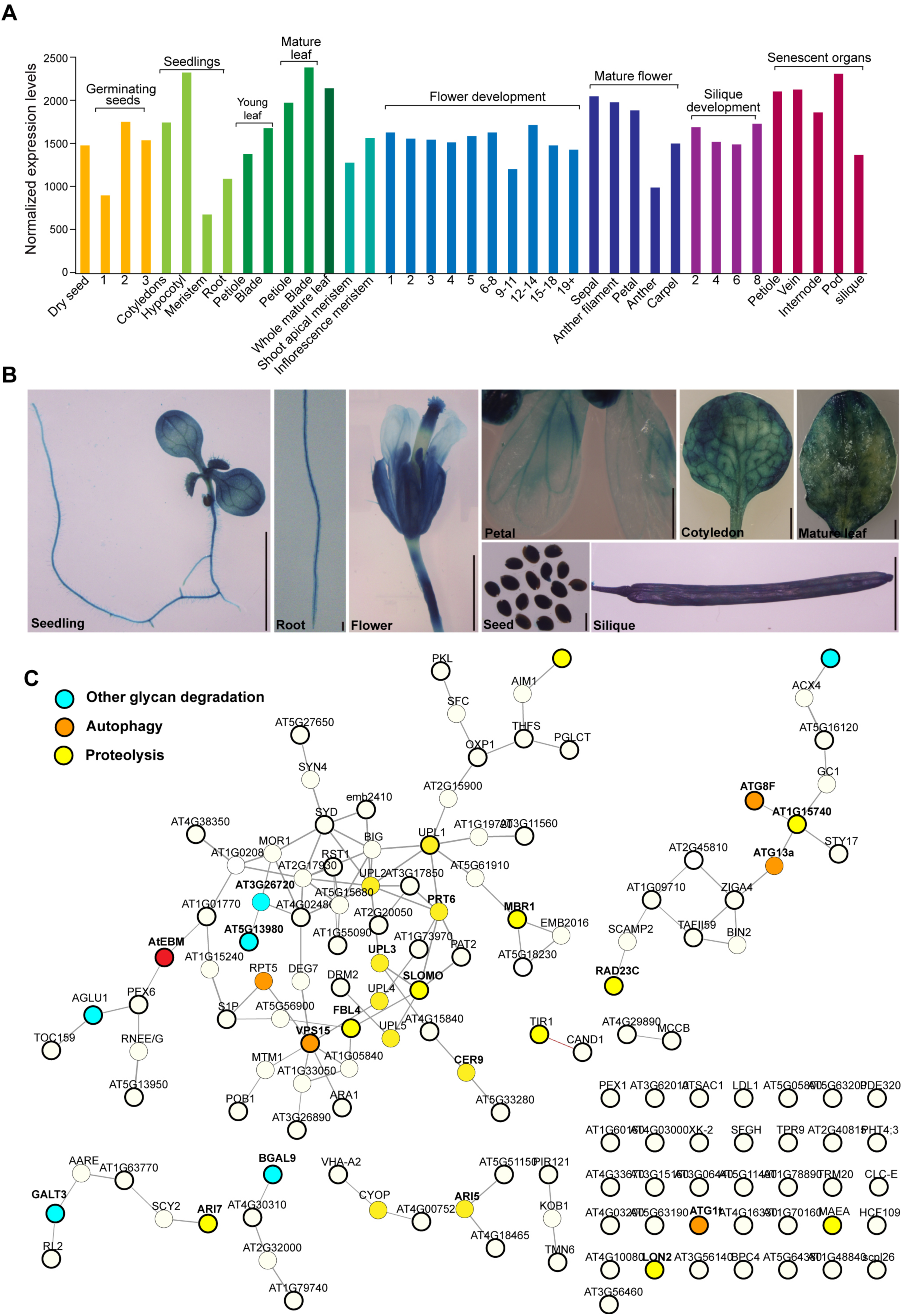
Expression patterns of the *AtEBM* gene. **(A)** *AtEBM* mRNA expression levels across plant development. Data were downloaded from TraVA (http://travadb.org/; Klepikova *et al*. 2016). Germinating seed samples were collected at 1–3 days after soaking. Flower development samples were collected at the moment of the anthesis of the first flower. The numbers represent the order of flower initiation. Silique development samples (seeds not removed) were harvested when the first silique reached 1 cm in length. The numbers represent the order of silique initiation. **(B)** GUS staining in *proAtEBM::GUS* transgenic plants. GUS staining in 7-day-old seedling, root, flower, petal, cotyledon, mature leaf, seed, and silique. Scale bar = 500 µm. **(C)** Co-expression network of AtEBM. The ATTED-II NetworkDrawer tool was used to construct the co-expression network using the first 100 genes associated with AtEBM as bait genes (thicker circles) to search the ATTED-II database (https://atted.jp/). The candidate gene AtEBM is highlighted in a red circle. Cyan, yellow, and orange circles indicate genes involved in other glycan degradation, proteolysis, and autophagy, respectively, which are significantly enriched based on Gene Ontology (GO) analysis.

**Supplemental Figure 6.**
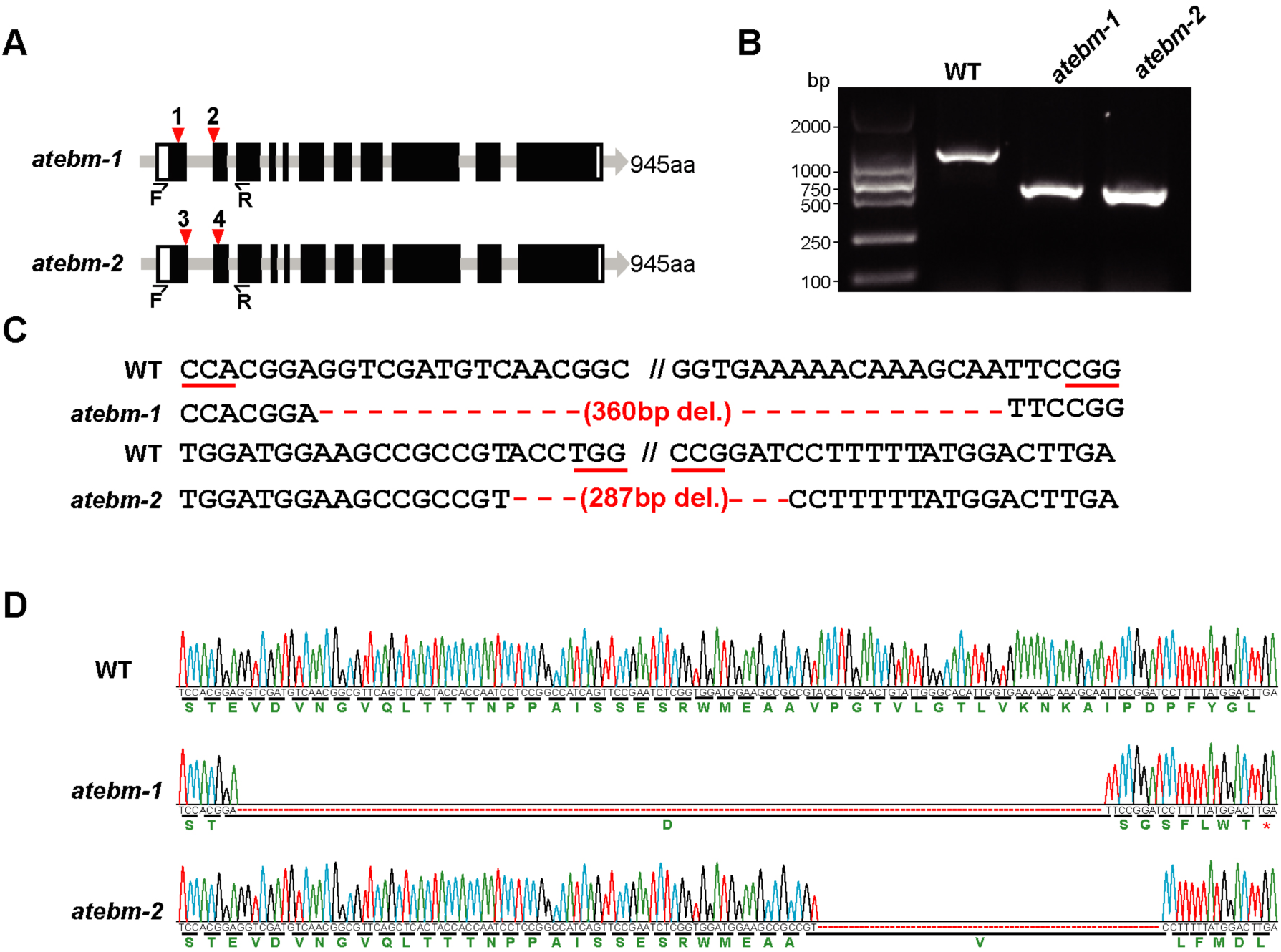
Genome editing of *Arabidopsis AtEBM* gene using CRISPR/Cas9 system. **(A)** Schematic diagram of *AtEBM* gene with four targeting sites indicated by red triangles. The black boxes indicate exons and the white boxes show untranslated regions. The half arrows indicate the primers used for genomic PCR in (B). **(B)** Genomic PCR analysis of the *atebm* deletion mutants. Total genomic DNA isolated from the wild type (WT), *atebm-1*, and *atebm-2* plants was subjected to PCR analysis using the primers indicated in (A). **(C)** Sequencing results of *atebm-1* and *atebm-2* alleles. Protospacer-adjacent motifs (PAMs) are underlined. **(D)** Sanger sequencing peaks of the *atebm-1* and *atebm-2* alleles. The chromatograms shown were generated by Sanger DNA sequencing of AtEBM cDNAs amplified from *atebm-1* and *atebm-2* mutants. The amino acid sequence translated from each trace is shown below the DNA sequence. Red asterisks denote stop codons, and dashes indicate gaps.

**Supplemental Figure 7.**
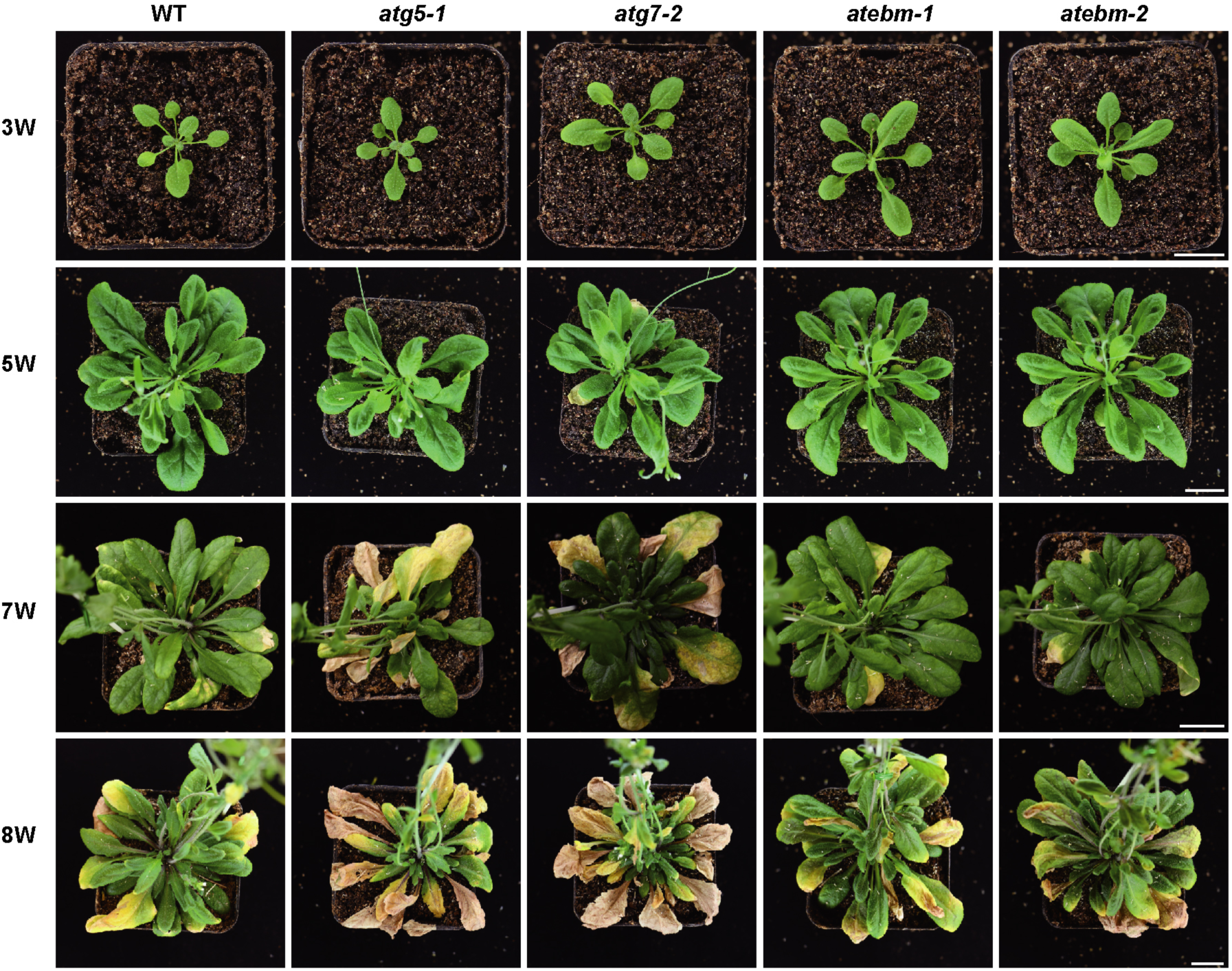
Loss of *AtEBM* does not affect plant development. The wild-type Col-0 (WT), autophagy mutants *atg5-1* and *atg7-2*, and *atebm* mutants *atebm-1* and *atebm-2* were grown at 22°C in soil under short-day conditions (12 hr light/12 hr dark). Photos were taken 3, 5, 7, and 8 weeks after germination. Bars = 2 cm.

**Supplemental Figure 8.**
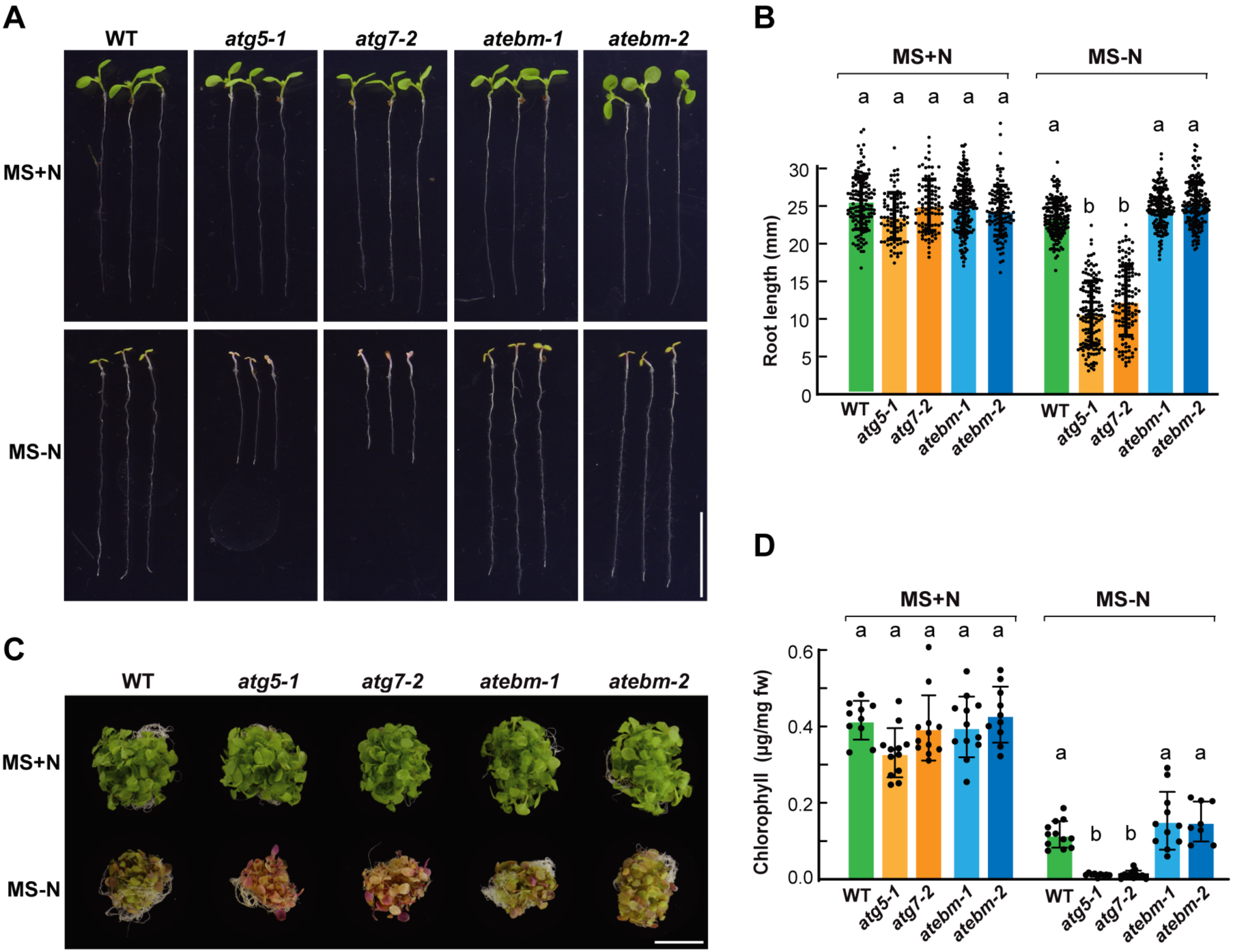
*atebm* mutants were insensitive to nitrogen starvation. **(A-B)** Primary root phenotype of *atebm* mutants in response to nitrogen deficiency. Seeds were germinated and grown vertically on MS medium with or without nitrogen under LD conditions for one week. Bar = 1 cm. (B) Quantification of the root length of plants shown in (A). Data are presented as means ± s.d. (*n* = 3 biological replicates, >100 seedlings per genotype in each independent experiment). **(C-D)** Leaf senescence phenotypes of *atebm* mutants in response to nitrogen starvation. Seeds were germinated in an MS liquid medium containing nitrogen and grown for one week under constant white light conditions, and then transferred to either a fresh MS (+N) or a nitrogen-deficient (–N) liquid medium for an additional week. Total chlorophyll content of the plants shown in (D). Data are presented as means ± s.d. (*n* = 3 biological replicates: 80-120 seedlings per genotype in each independent experiment).

**Supplemental Figure 9.**
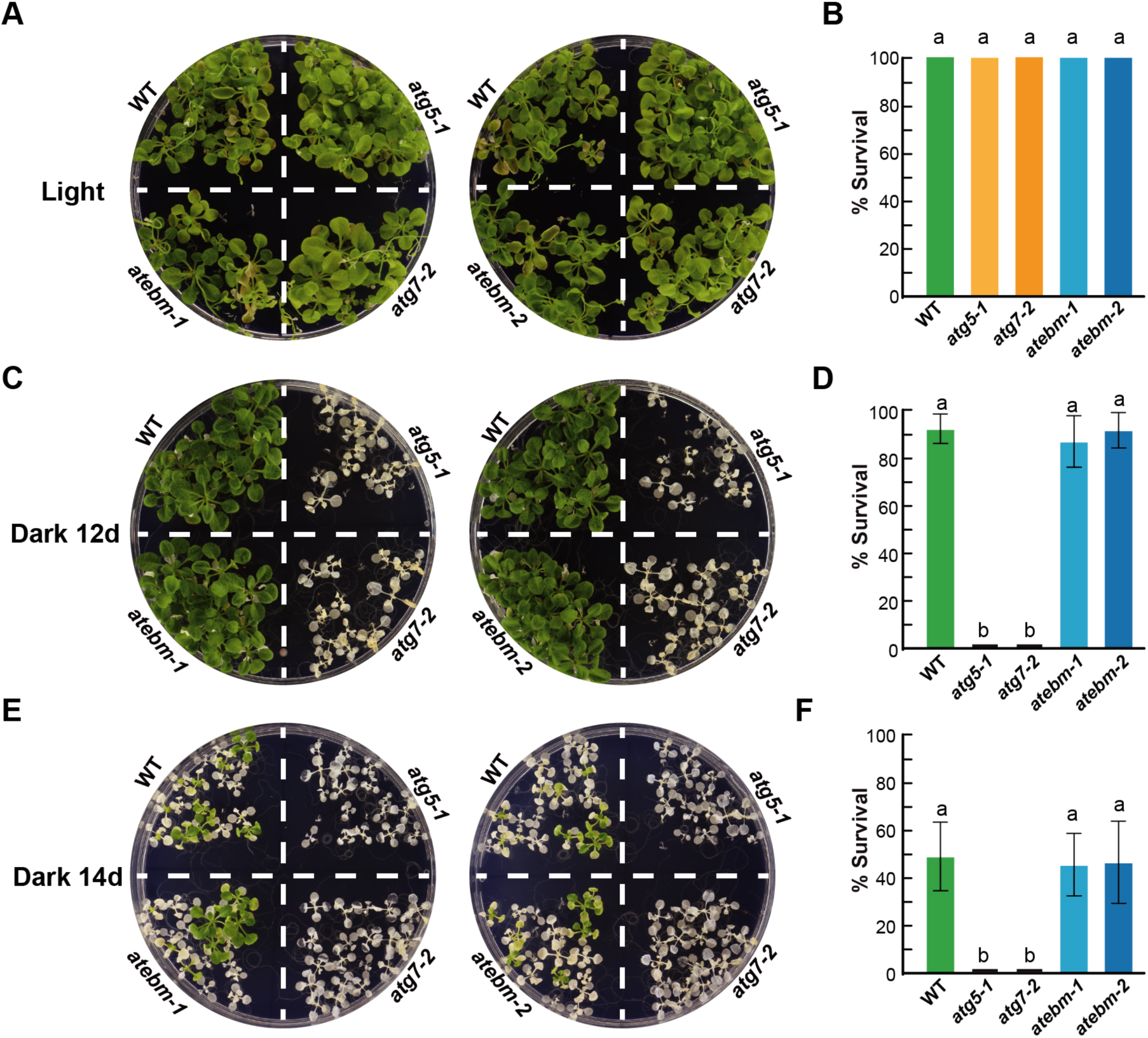
*atebm* mutants were insensitive to fixed-carbon starvation. The seedlings of wild-type Col-0 (WT), autophagy mutants *atg5-1* and *atg7-2*, *atebm* mutants *atebm-1* and *atebm-2* were grown under LD conditions on MS solid medium lacking Suc for two weeks, then transferred to darkness for 12 days (C) or 14 days (E), or remained under the same light condition (A). After the dark treatment, the stressed plants were transferred back to LD conditions for 12 days, after which the survival rates of the seedlings (B, D and F) were quantified to evaluate the effects of fixed-carbon starvation. Data are presented as means ± s.d. (*n* = 3 biological replicates; 60-120 seedlings per genotype in each independent experiment).

**Supplemental Figure 10.**
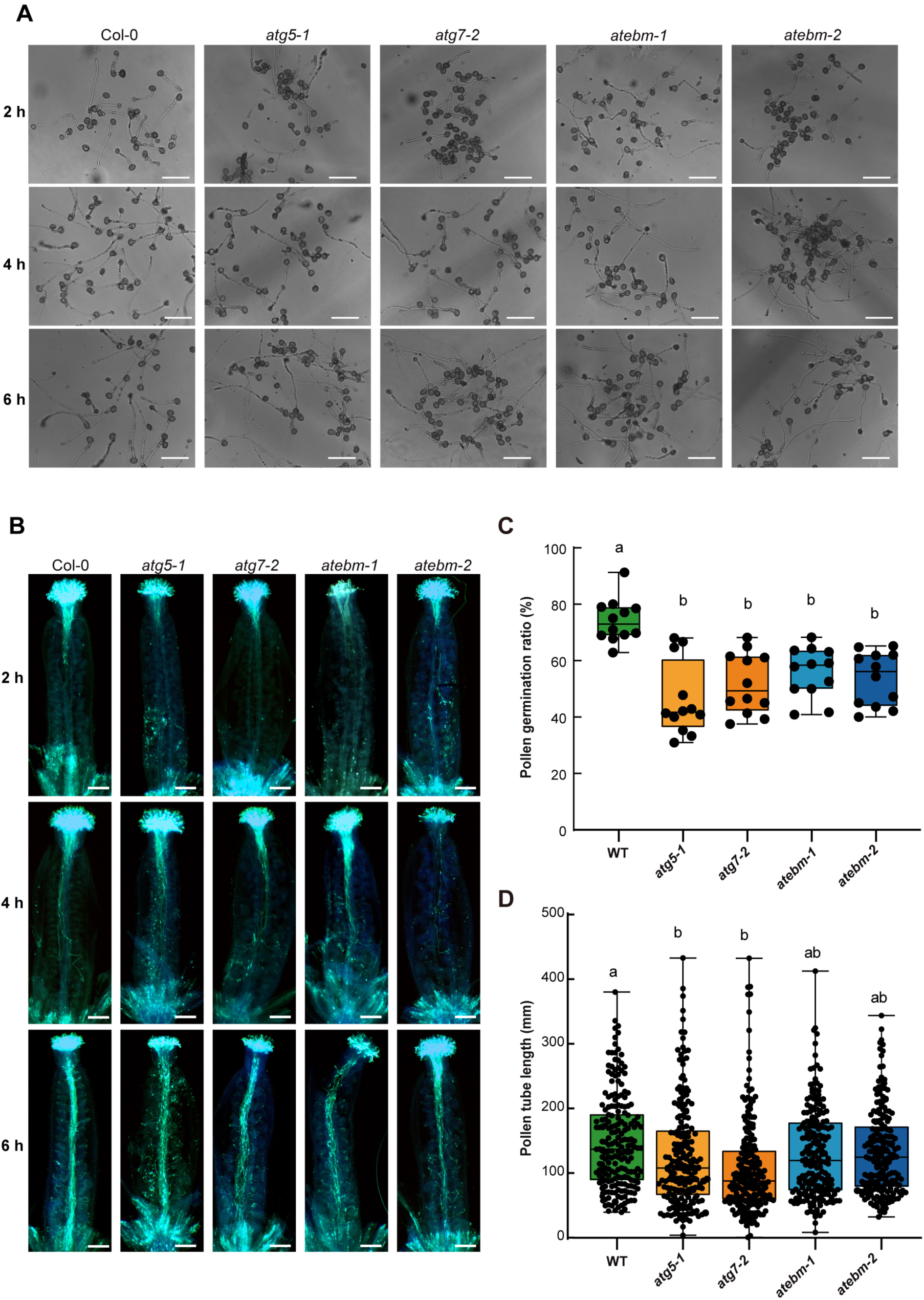
*In vitro* and *in vivo* germination of the pollens of *atebm* mutants. (A) *In vitro* pollen germination assays for the *atebm* mutants. Representative photomicrographs of pollen germinated on medium for 2, 4 and 6 hours. Bars = 10 μm. (B) *In vivo* pollen germination assay. Aniline blue staining of pollens after pollination for 2, 4, and 6 hours. Bars = 10 μm. (C) Quantitative analysis of pollen germination rate after 2 h. Different letters indicate significant differences (*p* < 0.05) as determined by two-way ANOVA followed by Tukey’s multiple comparison test. (D) Quantitative analysis of the pollen tube lengths of WT, *atg5-1, atg7-2*, and *atebm* mutants after 2 hours growth *in vitro* on a germination medium. Data are presented as means ± s.d. of three biological replicates.

